# Dissociable roles of thalamic nuclei in the refinement of reaches to spatial targets

**DOI:** 10.1101/2023.09.20.558560

**Authors:** LJ Sibener, AC Mosberger, TX Chen, VR Athalye, JM Murray, RM Costa

## Abstract

Reaches are complex movements that are critical for survival, and encompass the control of different aspects such as direction, speed, and endpoint precision. Complex movements have been postulated to be learned and controlled through distributed motor networks, of which the thalamus is a highly connected node. Still, the role of different thalamic circuits in learning and controlling specific aspects of reaches has not been investigated. We report dissociable roles of two distinct thalamic nuclei – the parafascicular (Pf) and ventroanterior/ventrolateral (VAL) nuclei – in the refinement of spatial target reaches in mice. Using 2-photon calcium imaging in a head-fixed joystick task where mice learned to reach to a target in space, we found that glutamatergic neurons in both areas were most active during reaches early in learning. Reach-related activity in both areas decreased late in learning, as movement direction was refined and reaches increased in accuracy. Furthermore, the population dynamics of Pf, but not VAL, covaried in different subspaces in early and late learning, but eventually stabilized in late learning. The neural activity in Pf, but not VAL, encoded the direction of reaches in early but not late learning. Accordingly, bilateral lesions of Pf before, but not after learning, strongly and specifically impaired the refinement of reach direction. VAL lesions did not impact direction refinement, but instead resulted in increased speed and target overshoot. Our findings provide new evidence that the thalamus is a critical motor node in the learning and control of reaching movements, with specific subnuclei controlling distinct aspects of the reach early in learning.

## INTRODUCTION

Skillfully reaching to targets in space is an important component of many actions, such as bringing a paint brush to a canvas. Reaches are learned and controlled through distributed motor networks, encompassing circuits such as the motor cortex^1–5^, basal ganglia^6,7^, cerebellum^8–10^, and brainstem^11–13^. The basal ganglia and cerebellar circuits have been seen as having complementary roles in motor learning and execution, with the basal ganglia more involved in control and reinforcement of specific movements^14–18^, and the cerebellum providing sensory feedback during ongoing actions to regulate the speed, smoothness, and endpoint precision of reaching actions^10,19^.

There is clear evidence that the thalamus, which is comprised of nuclei that receive inputs and send outputs to different motor centers, is a critical node in movement and motor learning^20,21^. Particularly, the thalamic parafascicular (Pf) and the ventroanterior/ventrolateral (VAL) nuclei are situated at the convergence of important motor centers. Pf receives inputs from basal ganglia output nuclei, the internal segment of the globus pallidus (GPi) and substantia nigra pars reticulata (SNr)^22,23^. Pf’s outputs include somatotopically organized glutamatergic projections to the striatum, in almost equal amount with the motor cortex^20,23,24^, and to the subthalamic nucleus (STN)^25^. VAL, on the other hand, receives excitatory driver-like inputs from deep cerebellar nuclei (DCN)^26–28^ and inhibitory inputs from the basal ganglia^26,28^, and outputs primarily targeting the deep and superficial layers of motor cortex^29,30^. Pf and VAL’s separate anatomical footholds in these distributed motor networks suggests that they may underlie different aspects of learning and movement. Indeed, there is evidence that Pf and VAL are important for learning and movement^25,31,32^. However, it is unknown what exact components of complex movements they encode and control, and how their roles evolve with learning.

Here, we investigated how Pf and VAL contribute to the control and refinement of reaches to a target in space, using a head-fixed joystick task that we developed^33^. We used 2-photon calcium imaging of glutamatergic neural populations in the forelimb regions of both areas across time, and uncovered that many neurons change activity during reaches early learning. This reach-related activity decreased as learning progressed. By tracking the same neurons over time, we uncovered that population dynamics in Pf changed between early and late learning, but were stable after learning. In contrast, VAL’s dynamics continuously shifted regardless of learning stage. Importantly, we show that neural activity in Pf, but not VAL, encodes the direction of reaches early in learning, but not after a target has been learned and reaches refined. In accordance, pre-learning genetically targeted lesions revealed that glutamatergic neurons in Pf were required to refine reach direction during learning. VAL lesions, on the other hand, did not impair the refinement of reach direction, but resulted in target overshoot and faster movements. Post-learning lesions of both areas did not affect the performance of learned reaches. These results demonstrate that Pf and VAL are critical nodes for motor behavior with distinct roles in forelimb reaching movements, and broadly that the thalamus contains separable movement governing circuits.

## RESULTS

### Mice explore and refine directional reaches to a spatial target

To study the roles of Pf and VAL in the learning and refinement of target reaches, we trained mice (n = 10) in our Spatial Target Task (STT)^33^ while we imaged their thalamic neural activity (**Figure 1A/B**). Head-fixed mice explored a 2-dimensional workspace by moving a joystick with their right forelimb from a set start location. Movements were rewarded when attempts entered a circular spatial target (hits) or aborted when the animal let go of the joystick or a maximum time (7.5 seconds) elapsed (misses) (**Figure 1C**). During pre-training, mice were rewarded for movements in the general forward direction, and mouse-specific targets 40° to the left and right of the mean direction were defined on the last day of pre-training (**Figure S1A/B**). After target definition, mice were trained in two behavioral blocks with one active target per block such that reaches in different directions were learned in each block (**Figure 1D/E**). The active target was changed from target 1 to target 2 the day after the animal reached a high-performance criterion on target 1 (>60% average hit ratio over three consecutive days), or a maximum number of days (53 days) had elapsed (**Figure S1C**). In order to compare results across animals with different block lengths, we selected five equidistant days per block for each animal (see Methods).

**Figure 1.**
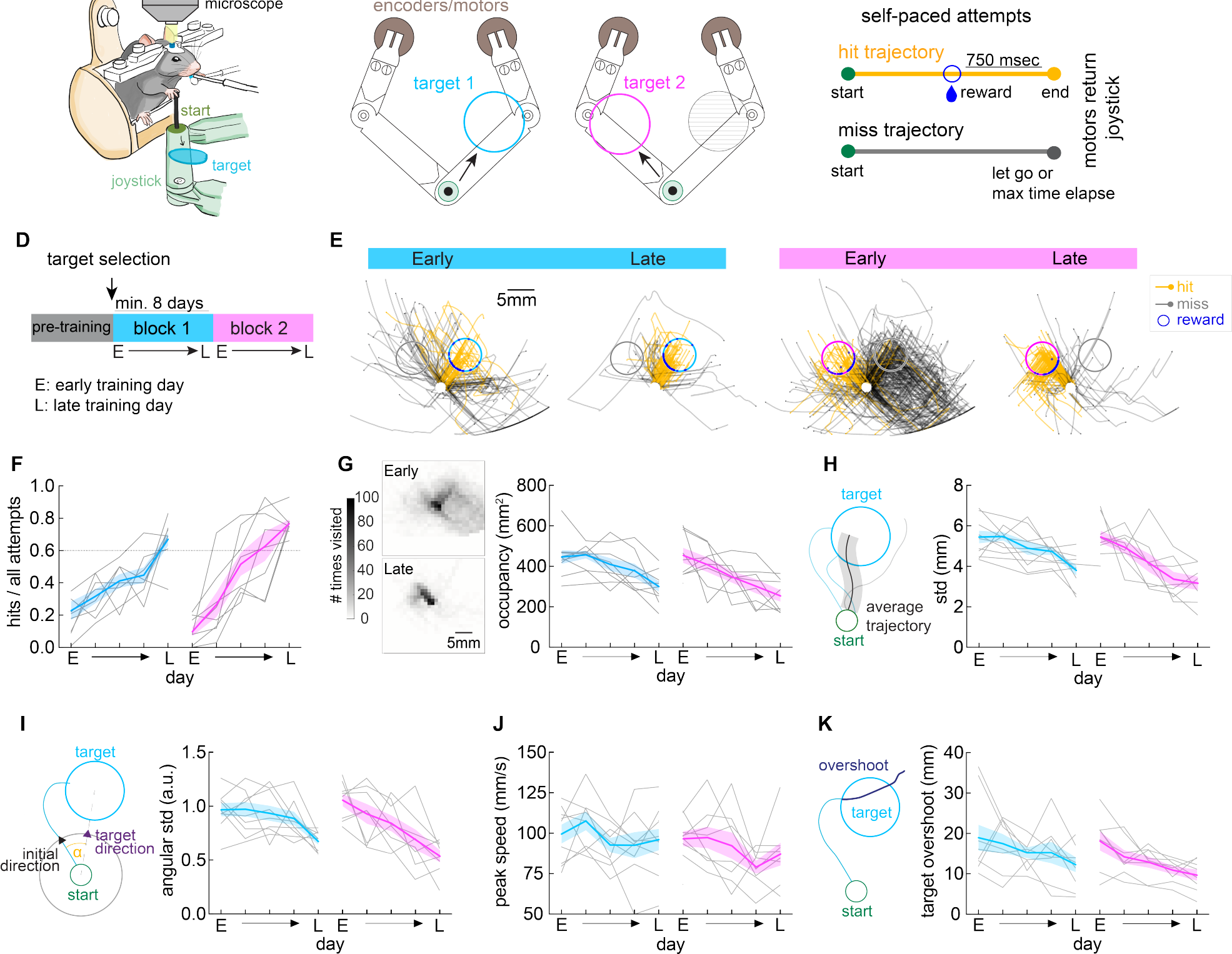
Mice explore and refine directional spatial target reaches. (A)Schematic of a head-fixed mouse performing the Spatial Target Task (STT) during imaging. (B) Left: top view schematic of the Selective Compliance Articulated Robot Arm (SCARA) joystick showing the start position (green circle) and example position of target 1 (blue circle). Right: same as left, example position of target 2 (magenta circle) and previous target 1 (gray dashed area). (C) Classification of trajectories from self-paced joystick movements. Hit trajectories (yellow line), and miss trajectories (gray line), started with the mouse moving the joystick out of the start position (green circle). Hit trajectories were rewarded upon target hit and ended by motors returning the joystick to the start 750 msec later. Miss trajectories were aborted by motors returning the joystick to the start if the animal let go of the joystick or the attempt timed out (after 7.5 sec). (D) Schematic showing the phases of the training protocol: during ‘pre-training’ animals were rewarded first for touching the joystick, then for pushing it in a forward direction. During blocks 1 and 2, attempts that entered targets 1 or 2 were rewarded, respectively. (E) Left: representative example trajectories on early and late days of block 1 with rewarded target (blue circle) and non-rewarding target (gray circle). Right: same for block 2 with rewarded target (magenta circle) and non-rewarding target (gray circle). Hit trajectories (yellow) that enter the rewarded target and miss trajectories (grey) are plotted in the two-dimensional workspace. Point of entering target and reward delivery is shown as small dark blue circle. (F) Hit ratio on 5 selected days per block from early training day (E) to late training day (L) and 3 equidistant days in between for each animal (mixed-effects model: day in block: *p < 0.001, day × mouse: p = 0.037) (G) Left: representative example heatmap of workspace occupancy of all trajectories of an early and late session (# times a bin is visited). Right: occupancy of workspace (mm^2^) (mixed-effects model: day in block: *p < 0.001, mouse: *p = 0.039) (H) Variability of mean trajectory from all movements (averaged std along the length of the trajectory, mm) (mixed-effects model: day in block: *p < 0.001, mouse: *p = 0.006). (I) Variability of initial direction of movement (angular std, a.u.) (mixed-effects model: day in block: *p = 0.002, mouse: *p = 0.002). (J) Peak speed of trajectories (mm/sec) (mixed-effects model: day in block: *p = 0.013, mouse: *p = 0.002) (K) Average target overshoot (mm) from target entry for all hit trajectories (mixed-effects model: day in block: *p < 0.001, mouse: *p = 0.016). (F-K) Mean ± SEM is shown in thick color lines and shaded bounds (block 1: blue, block 2: magenta), single animals are shown in gray lines (n = 10).

Mice missed the target in most attempts early in each block and gradually increased their performance to reach high hit ratios late in the block (**Figure 1F**). As animals learned to hit the rewarded target area, their reach trajectories occupied less of the workspace (**Figure 1G**), and the variability of their mean trajectory significantly decreased (**Figure 1H**). To investigate if mice refined the initial direction of reaching movements, we measured the variability of the initial movement vector angle using the angular std^34^, and found that it significantly decreased within each block (**Figure 1I**). In addition to learning to reach in the correct direction, animals also decreased the peak speed of their reaches (**Figure 1J**), suggesting that they slowed down to increase their chance of hitting the target. In line with this finding, we also found that mice significantly reduced the target overshoot pathlength with learning of each target (**Figure 1K**).

These findings suggest that as mice learn to reach the rewarded targets, they refined the initial direction of movement and reduced peak speed, while increasing targeting precision. Furthermore, on the first day of block 2 when the new target was rewarded, even though mice initially perseverated with movements to the previously rewarded target (**Figure S1D**), they promptly increased the variability of their trajectory and initial reach direction (**Figure 1H/I**). This change in behavior indicates an increase in exploration to discover the new target direction.

### Pf_FL_ and VAL_FL_ neural populations are most active during reaches early in learning

To investigate the role of thalamic areas in the learning and performance of forelimb reaches, we mapped and targeted the forelimb-related thalamus for our imaging experiments. Previous studies have detailed the somatotopy of primary motor cortex (M1) and identified a consistent caudal forelimb area (CFA) and rostral forelimb area (RFA)^35^. Similarly, the dorsal striatum has been shown to contain distinct areas that receive input from sensorimotor cortex according to their somatotopy^36,37^, which identified a dorsolateral striatal forelimb-related area (DLS_FL_). Strongly projecting to these major sensorimotor regions, Pf and VAL have been demonstrated to have a consistent topographic organization as well^20,38,39^. To specifically map the forelimb-related thalamus we injected the retrograde tracer cholera toxin subunit B (CTB) into DLS_FL_ and the CFA of M1 in wildtype mice (**Figure S2A/B and E/F**). We serially processed and reconstructed the brains using the BrainJ^40^ pipeline and aligned them to the updated Allen Brain Atlas (ABA)^41^. We found that Pf and VAL were the most strongly labeled thalamic nuclei when CTB was injected into DLS_FL_ and CFA, respectively (**Figure S2C/G**). Using these data, we delineated the forelimb-related thalamic areas (Pf_FL_ and VAL_FL_) by reconstructing the regions with detected cell bodies and integrated those volumes as new areas in the ABA (**Figure S2D/H**).

To measure the activity of glutamatergic cells in Pf_FL_ and VAL_FL_ during learning and performance of targeted reaches, we injected AAV5 expressing GCamp6f under a CamKII promoter into these respective areas, followed by implantation of a gradient index (GRIN) lens, and recorded single cell calcium activity of neurons using a 2-photon microscope throughout training of the STT in BL6 mice (**Figure 2A**). Cells were segmented and calcium transients extracted using Suite2p^42^ (**Figure 2B**). Individual cell traces were detrended and z-scored across daily sessions for all further analysis (**Figure 2C**). Imaging yielded a stable number of cells across several weeks of behavioral training (Pf: average neurons / animal / day 34 ± 3, VAL: average neurons / animal / day 33 ± 4) (**Figure S3**). In order to investigate whether neural activity in Pf_FL_ and VAL_FL_ was modulated during reaching movements, we aligned the neural activity to movement initiation (leaving start position) (**Figure 2C**) and averaged their activity across trials (**Figure 2D)**. In both thalamic areas, cells displayed diverse responses, including increasing, decreasing, or maintaining their average activity around the movement initiation. To test whether their activity was most modulated before, during, or after the target reach, we compared the ratio of modulated cells during three time windows around movement on the early day of block 1: pre-movement, -300-0 ms before movement initiation; during movement, 0-300 ms after leaving the start position; and after target hit, 0-300 ms following target hit (**Figure 2E)**. Cells were classified as significantly modulated if their trial-averaged activity within the defined time window was above or below 3x the SD of that cell’s baseline (calculated outside of the three windows). For Pf_FL_ cell populations, a significantly larger proportion of cells were modulated during the reach than before reach start, or after target hit (**Figure 2F, top**). VAL_FL_ populations were also strongly modulated during the movement window, but not significantly more than the other time windows (**Figure 2F, bottom**).

**Figure 2.**
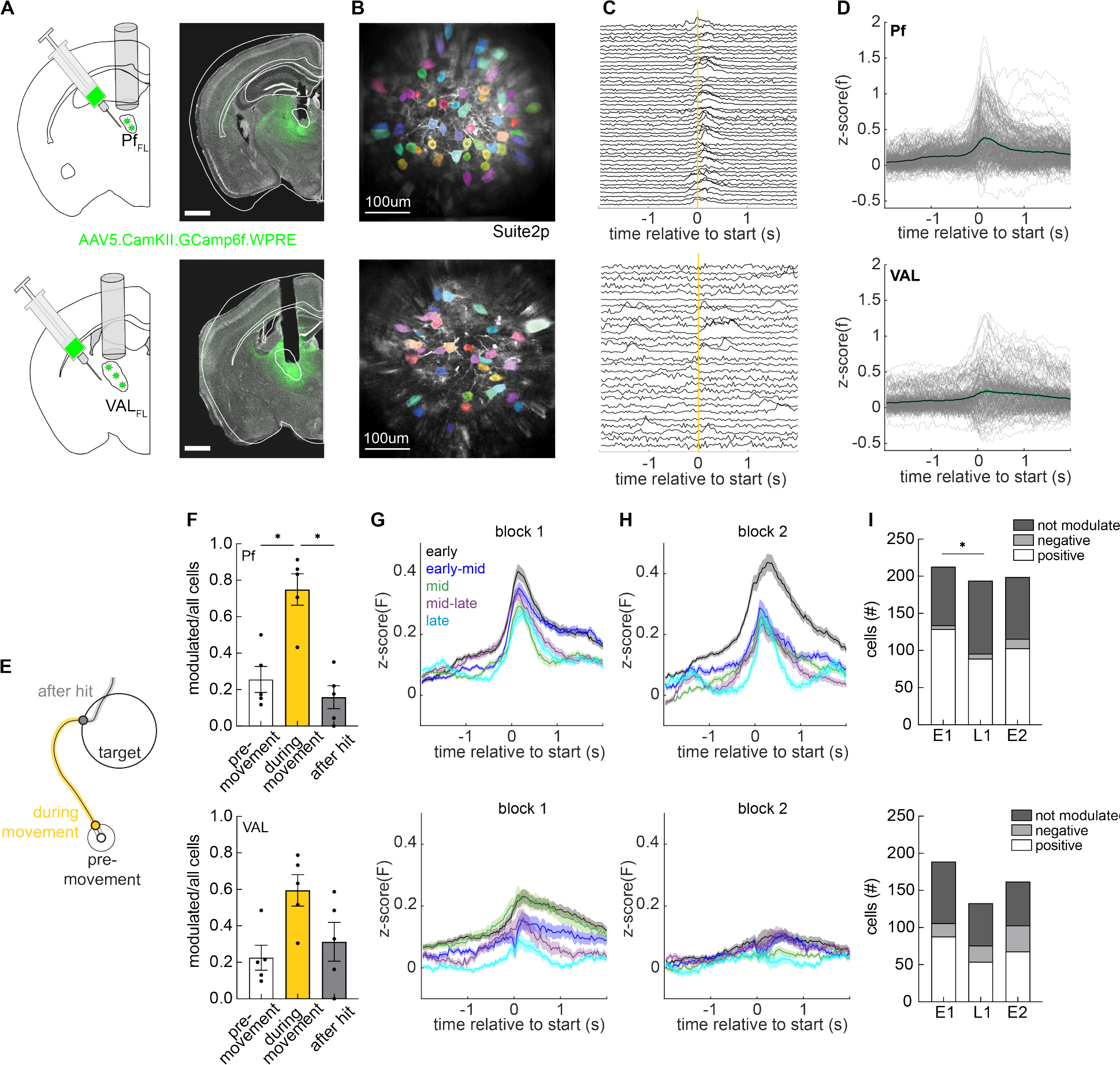
Pf_FL_ and VAL_FL_ neural populations are most active during reaches early in learning. (A)Top: schematic and representative histology of viral targeting and GRIN lens implantation for calcium imaging in Pf_FL_. Bottom: same for VAL_FL_. Scale bar represents 1 mm. (B)Top: example average fluorescence of the imaging field of view with detected ROI masks overlayed for Pf_FL_. Bottom: same for VAL_FL_. Scale represents 100 μm. (C)Top: example detrended and z-scored calcium traces of cells aligned to a reach start (yellow line) for Pf_FL_. Bottom: same for VAL_FL_. (D)Top: trial-averaged z-scored fluorescence aligned to reach start for Pf_FL_ cells on example day. Mean ± SEM (black trace and green shaded bounds). Individual cell trial-averages (gray); n = 213 neurons, 5 mice. Bottom: same for VAL_FL_; n = 189 neurons, 5 mice. (E)Schematic of the three time windows of a reach; pre-movement (white), during movement (yellow), after hit (gray). (F)Top: proportion of significantly modulated cells on early day of block 1 during the three time windows depicted in (E) for Pf_FL_ (repeated measures one-way ANOVA: *p = 0.006; Šídák’s multiple comparisons test: pre-movement vs during movement: *p = 0.002; during movement vs after hit: *p = 0.028, n = 5 mice). Bottom: same for VAL_FL_. (repeated measures one-way ANOVA: p = 0.078, n = 5 mice.) (G)Top: trial-average z-scored fluorescence of all cells on 5 sessions from block 1; early (black), early-middle (blue), middle (green), middle-late (purple), late (cyan) for Pf_FL_ (Kruskal-Wallis test, population activity at start for early vs late: *p < 0.001). Bottom: same for VAL_FL_ (Kruskal-Wallis test, population activity at start for early vs late: *p < 0.001). Data shown as mean ± SEM. (H)Same as (G), but over 5 session from block 2; Top: Pf_FL_ (Kruskal-Wallis test, population activity at start for early vs late: *p < 0.001). Bottom: same for VAL_FL_ (Kruskal-Wallis test, population activity at start for early vs late: p = 0.472). (I) Top: number of Pf_FL_ cells that are positively modulated (white), negatively modulated (light gray), or not modulated (dark gray), during the movement window on the early day of block 1 (E1), late day of block 1 (L1), and early day of block 2 (E2) (Chi-squared test, early block 1 vs late block 1: *p = 0.012; late block 1 vs early block 2: p = 0.135). Bottom: same for VAL_FL_ (Chi-squared test, early block 1 vs late block 1: p = 0.154; late block 1 vs early block 2: p = 0.412). VAL_FL_ showed significantly more negatively modulated cells than Pf_FL_ (Chi-square test, Pf vs VAL early block 1: *p = 0.001; late block 1: *p < 0.001; early block 2: *p < 0.001).

We next asked whether this reach-related activity changed over the course of learning in either thalamic area. To this end, we created trial-averaged, population-averaged activity traces for all of the five selected days of each block. In both blocks, we found that the overall activity in Pf_FL_ was high early in the block, but significantly decreased with learning as the reaches refined (**Figure 2G/H, top**). Activity in VAL_FL_ also decreased with learning in the first block, but did not return to a high magnitude early in block 2 and was low throughout the second block (**Figure 2G/H, bottom)**. This decrease in average activity over learning could be explained by a change in the relative numbers of positively and negatively modulated cells. We found that the relative numbers of modulated cells in Pf_FL_ changed from early to late days in block 1 (**Figure 2I, top**). This indicates that more cells in Pf_FL_ were positively modulated early in training than late in training. VAL_FL_ modulated cell ratios remained relatively consistent throughout training (**Figure 2I, bottom**).

This analysis also revealed that there was a larger ratio of negatively modulated cells in VAL_FL_ compared to Pf_FL_ across all analyzed days (**Figure 2I**), explaining the overall lower average activity in VAL_FL_. These results provide evidence that in our task Pf_FL_ and VAL_FL_ cells are mostly active during the execution of forelimb reaches, and that the magnitude of the population activity decreases with learning as fewer cells were positively modulated during the reach as training progressed. Interestingly, PfFL’s population activity increased again after target switch, suggesting that it is not just habituation but that Pf neurons were indeed more active early in learning.

We next asked if the observed changes in population activity were due to different cells being engaged at different stages of learning, or the same cells modulating their activity differently across learning stages. To investigate this, we repeated the previous analysis using only cells that were matched across the late day of block 1 and the early day of block 2 (**Figure 3A**). By focusing on this particular point in behavior, we captured two days close in time but at different learning stages; a day with refined reaching movements and high performance (late) and a day with exploratory variable reaches with low performance (early). To match the cells across days, we used non-rigid motion correction to align the imaging fields of view from the selected days and matched the segmented cells (**Figure S4)** (Pf, n = 116 cells, 4 mice; VAL, n = 86 cells, 5 mice). We found that the reach-related population activity of matched cells in Pf_FL_ was still significantly higher on the early day of block 2 compared to the late day of block 1 (**Figure 3B, left**). Similar to the results from the entire population, this was not the case for the matched VAL_FL_ populations (**Figure 3B, right**). To control for baseline population changes across days, we repeated the analysis for two days late in block 1. This captures another pair of days close in time, but now with similar behavioral profiles (**Figure S5A**). Matched cell population activity did not change across these days, when reaches had already refined and behavioral variability was low (Pf_FL_, n = 108 cells, 4 mice; VAL_FL_, n = 72 cells, 4 mice) (**Figure S5B**).

**Figure 3.**
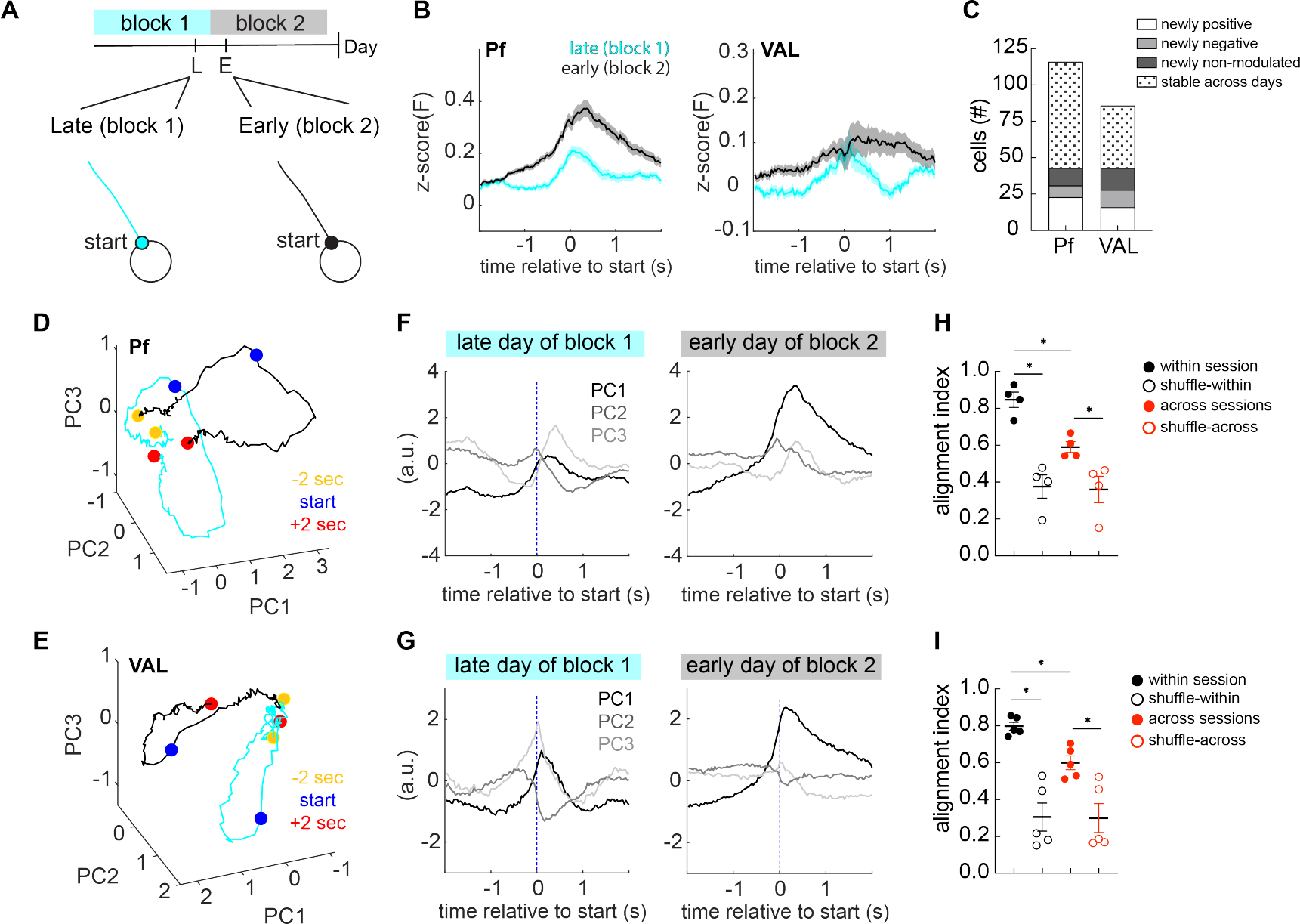
Low dimensional population activity alignment changes between early and late learning. (A)Schematic showing the selected days for cell matching; late day of block 1 (L) and early day of block 2 (E) and schematic trajectory around movement start. (B)Left: trial-averaged fluorescence aligned to movement start for matched cells on late block 1 (cyan) and early block 2 (black) for Pf_FL_ (n = 116 cells, 4 mice; Wilcoxon matched-pairs signed rank test: *p < 0.0001). Right: same for VAL_FL_ (n = 86 cells, 5 mice; Wilcoxon matched-pairs signed rank test: p = 0.577) Mean ± SEM (thick colored line with shaded bounds). (C)For Pf_FL_ and VAL_FL_ matched cells, stable and changing cell modulation on the early day of block 2 compared to the late day of block 1. Cells became newly positively modulated (white), newly negatively modulated (light gray), newly non-modulated (dark gray), or were stable across days (dotted). (D) Neural activity of Pf_FL_ around movement start projected into top 3 PC space for late day of block 1 (cyan trace) and early day for block 2 (black trace). -2 seconds before movement onset (green circle), movement start (blue circle), and +2 seconds after movement start (red circle). (E) Same as (D), for VAL_FL_ matched neuronal populations. (F) Same neural activity as in (D) projected onto each of the top 3 PCs, plotted against time, and aligned to movement start (dashed blue line). (G) Same as (F), for VAL_FL_ matched neuronal populations. (H) For Pf_FL_, average alignment of subspaces occupied by top 3 PCs within (filled red) and across (filled black) sessions with matched cells (same sessions as D/E). Alignment within sessions is significantly higher than across, and controls (open circles) (2-way repeated measures ANOVA: Day of alignment: *p < 0.001; shuffle: *p = 0.006; Day of alignment x shuffle: *p = 0.006; Šídák’s multiple comparison: within vs across: *p = 0.004; within vs within-shuffle: *p < 0.001; across vs shuffle-across: *p = 0.005). Data shown as mean ± SEM. (I) Same as (H), for VAL_FL_. Alignment within sessions is significantly higher than across, and shuffle-controls (2-way repeated measures ANOVA: Day of alignment: *p = 0.011; shuffle: *p = 0.007; Day of alignment x shuffle: *p = 0.005; Šídák’s multiple comparison, within vs across: *p = 0.003; within vs within-shuffle: *p < 0.001; across vs shuffle-across: *p = 0.001).

Analyzing again the modulation of the matched cells across the transition from block 1 and block 2, we discovered that nearly 20% of Pf_FL_ matched cells changed their activity during movement and became newly positively modulated on the early day of the new target, while 63% of the population’s modulations were stable. This indicates that the increased population activity on the early day of block 1 was driven at least partially by a subpopulation of newly upmodulated cells (**Figure 3C**). The same analysis across the two late days of block 1 showed the largest proportion of cell modulation changes in Pf_FL_ came from 18% of cells becoming newly non-modulated, while 69% of the population’s modulations were stable. (**Figure S5C**). Overall, these results suggest that more Pf_FL_ neurons are active specifically during the early stages of learning a new target, and overall population activity reduced during reaches later in learning after movements were more precise. In comparison, we found that 14% of VAL_FL_ matched cells became newly negatively modulated across the transition from block 1 and block 2 (**Figure 3C**), while less than 2% of matched cells became newly negatively modulated across two late days in block 1 (**Figure S5C**). At the same time, 50% and 58% of VAL_FL_ cell modulations remained the same across late block 1 to early block 2, and two late days in block 2, respectively. These results suggest that cells in VAL_FL_ were changing their movement modulation across days, and that the increase of negatively modulated cells early in block 2 may have contributed to the low population average activity.

### The alignment of population dynamics changes between early and late learning

We next asked whether these activity changes on population and single cell level affected the low-dimensional neural population dynamics, which would suggest that they affect underlying computations in the area. To find the dominant patterns of covariance across neurons we applied principal component analysis (PCA) on the trial-averaged activity of the same matched cells around the time of movement start. In order to identify the principal components, or the dimensions of population activity that captured the most variance, across both days a single PCA was run on the concatenated time windows from late and early days. First, we visualized the population activity of each day in the top 3 PC-space. Movement-locked population activity appeared to occupy different dimensions in the early and late day within the space spanned by the first 3 PCs (explaining 73.5% and 74.2% of variance for Pf_FL_ and VAL_FL_, respectively) for both Pf_FL_ (**Figure 3D/F**) and VAL_FL_ (**Figure 3E/G**), with large changes along PC1 on the early day of block 2. In comparison, the activity projected onto the top 3 PCs appeared to traverse through PC space more similarly between the two days late in block 1 (**Figure S5D/F and E/G**).

To test if the dominant dimensions of neural covariance changed when the new target was rewarded, we calculated the principal components for each animal and day individually, and compared their alignment between split sessions within the same day versus across different days. We found that the dimensions containing the most variance were aligned within the same day, but these were significantly less aligned across late block 1 and early block 2 days for both Pf_FL_ and VAL_FL_ **(Figure 3H/I**). We again used the same analysis across two days late in block 1 to control for daily changes late in learning. The alignment indices within and across those sessions were not different for Pf_FL_ on the same target (**Figure S5H**), but alignment was lower across days for VAL_FL_ (**Figure S5I)**. Seeing that VAL_FL_ alignment indices were significantly different across both conditions (late block 1 – early block 2 and two days in early block 2), it may be that VAL_FL_’s neural activity continued to change the dimension of neural covariance each day regardless of learning stage. This is in stark contrast to Pf_FL_, which did not change the dominant dimensions of covariance late in learning. Further, we found that alignment within a session was reduced when neuron identity was shuffled (**Figure 3H/I)**. This result emphasizes that individual neurons had distinct activity profiles during movement; if all neurons shared a similar activity profile, then shuffling neuron identity would result in similar alignment. Additionally, this result shows that alignment across session was still greater than the chance level given by preserving temporal activity but randomizing its assignment to individual neurons. In summary, by tracking the same neurons across multiple days of training, we found that neural population dynamics of Pf_FL_ occupied different dimensions during early vs. late learning of directional reaches, which contrasted with the similar dimensions occupied during multiple days of stable performance in late learning, whereas VAL_FL_ occupied different dimensions each day even after learning.

### Pf_FL_neural activity predicts reach direction early in learning

Our analysis so far revealed that thalamic neuronal activity was strongly engaged during reaching movements, and this activity decreased with learning. For Pf_FL_, this activity increased again when the mouse learned a new target, with many cells becoming newly upmodulated, and neural population dynamics occupying different dimensions. To investigate whether this neural activity could be encoding the directional reaching movements, we trained linear Ridge regression models to decode the initial reach direction from the neural activity during the reach start (0-167 msec after start) (**Figure 4**). Regression models were trained to predict the sine and cosine of the initial vector angle on held out test trials (70%/30% train/test split, 10 models per session) (**Figure 4A-4D**), and the average coefficient of determination (R^2^) between the true direction and the predicted direction was used to compare model accuracy between animals and days. Separate models were trained on single sessions from early and late days in block 1 for all animals. The initial direction was inherently more variable on early days than late days, and we found that across all sessions model accuracy correlated with variability in the dependent variable. To compare early and late days, we thus subsampled the trials of all days to match the lowest reach direction variability of the dataset. Comparing the model accuracy between groups, we found that Pf_FL_ neural activity could predict the direction of movement during early learning significantly better than VAL_FL_ neural activity (**Figure 4E**). Additionally, this directional information could only be decoded from Pf_FL_ early in training, as we were unable to predict the direction of movement on late training days (**Figure 4E**). Models in which the trial identity was shuffled in the train dataset performed poorly on all days (**Figure 4E**). These results show that Pf’s movement-related activity early in learning contains directional information, while VAL’s does so to a lesser degree. This directional information was not present when the reaching movement had been refined, but returned once a new target was rewarded (data not shown).

**Figure 4.**
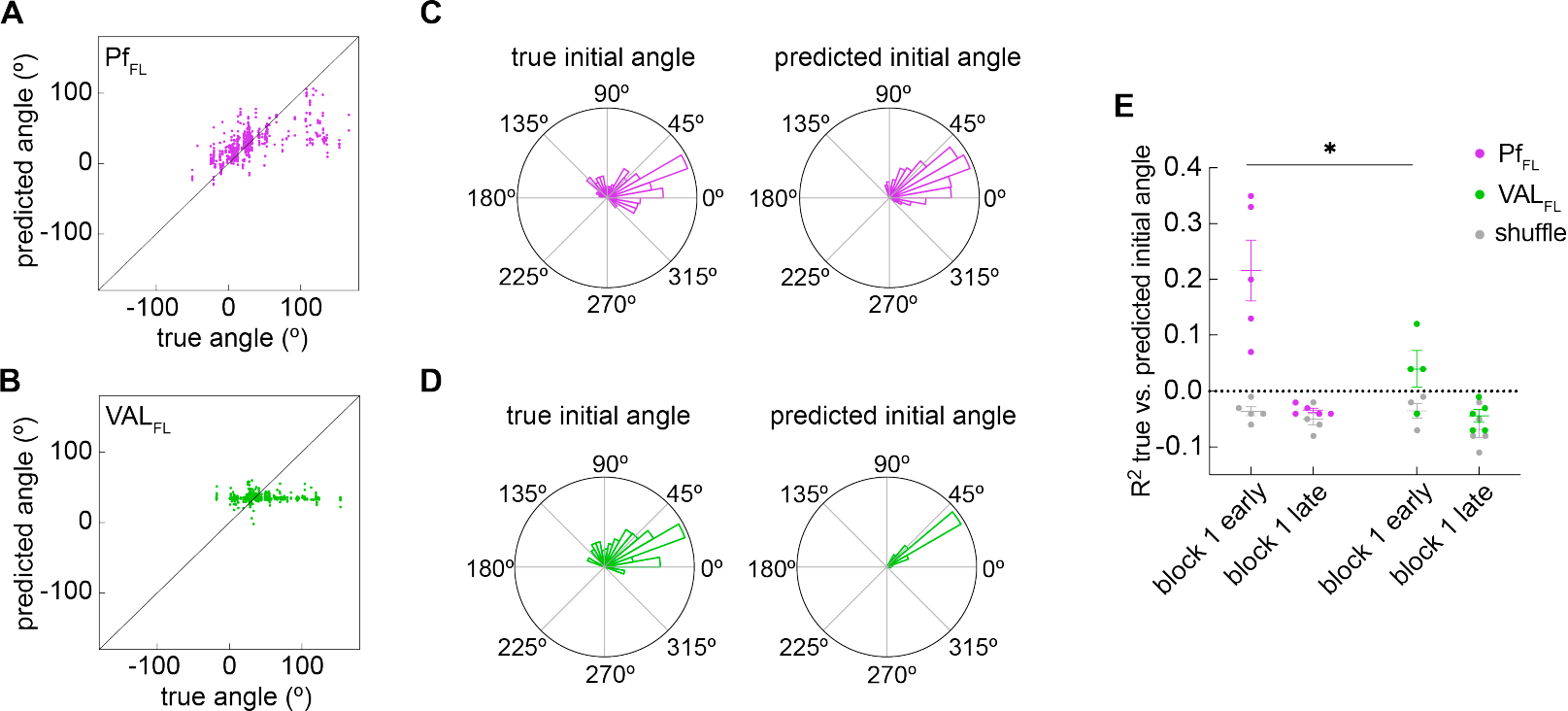
Pf_FL_ neural activity predicts reach direction early, but not late, in learning. (A)Predicted initial vector direction of combined held-out test data from 10 models trained on Pf_FL_ neural activity using different train/test splits, plotted against the corresponding true initial vector direction. Models trained on an early session in block 1. (B)Same as (A), for VAL_FL_ animal. (C)Circular histogram showing the true (left) and predicted (right) initial vector directions from Pf_FL_ animal data in (A) (D)Circular histogram showing the true (left) and predicted (right) initial vector directions from VAL_FL_ animal data in (B) (E)Averaged coefficients of determination (R^2^s) of model predictions for Pf_FL_ (purple), VAL_FL_ (green), and models trained on shuffled vector data (gray) on the early and late day of block 1. (Mixed-effects model: area (Pf or VAL), *p = 0.012; Šídák’s multiple comparisons test, Pf block 1 early vs VAL block 1 early, *p = 0.004).

### Ablation of Pf_FL_ but not VAL_FL_ cells before learning impairs the refinement of reach direction

Based on the results above, we hypothesize that Pf_FL_’s reach-related activity early in training, which predicted reach direction (**Figures 2 and 4**), is needed to learn and refine the direction of target reaches. To test this, we performed pre-learning lesions of Pf_FL_ and VAL_FL_. We selectively ablated glutamatergic cells by bilaterally injecting AAV5-flex-taCasp3-TEVp in the Pf_FL_ or VAL_FL_ of VGlut2-Cre mice and WT littermates (**Figure 5A**) (Pf_pre-lesion_, n=10; Pf_pre-control_, n=10; VAL_pre-lesion_, n=8; VAL_pre-control_, n=10), which induced cell death through apoptosis^43^. Lesions were confirmed by histological staining against the neuronal cell marker NeuroTrace (**Figure 5B-E**). Lesioned animals were left with large areas devoid of healthy NeuroTrace staining, with clear morphological differences to the WT control animals. Reconstruction of the 3D volumes of the lesions showed that the 34.33 *±* 16% of Pf_FL_ was lesioned, and 12.20 *±* 4% of VAL_FL_ (**Figure 5F, Figure S6A/B**). Even though our lesions left parts of Pf_FL_ and VAL_FL_ intact, we confirmed that Pf and VAL were the primary targets of the lesions over neighboring thalamic nuclei (**Figure S6C/D**). The percent of Pf_FL_ and VAL_FL_ lesioned did not correlate with overall performance (**Figure S6E/F**).

**Figure 5.**
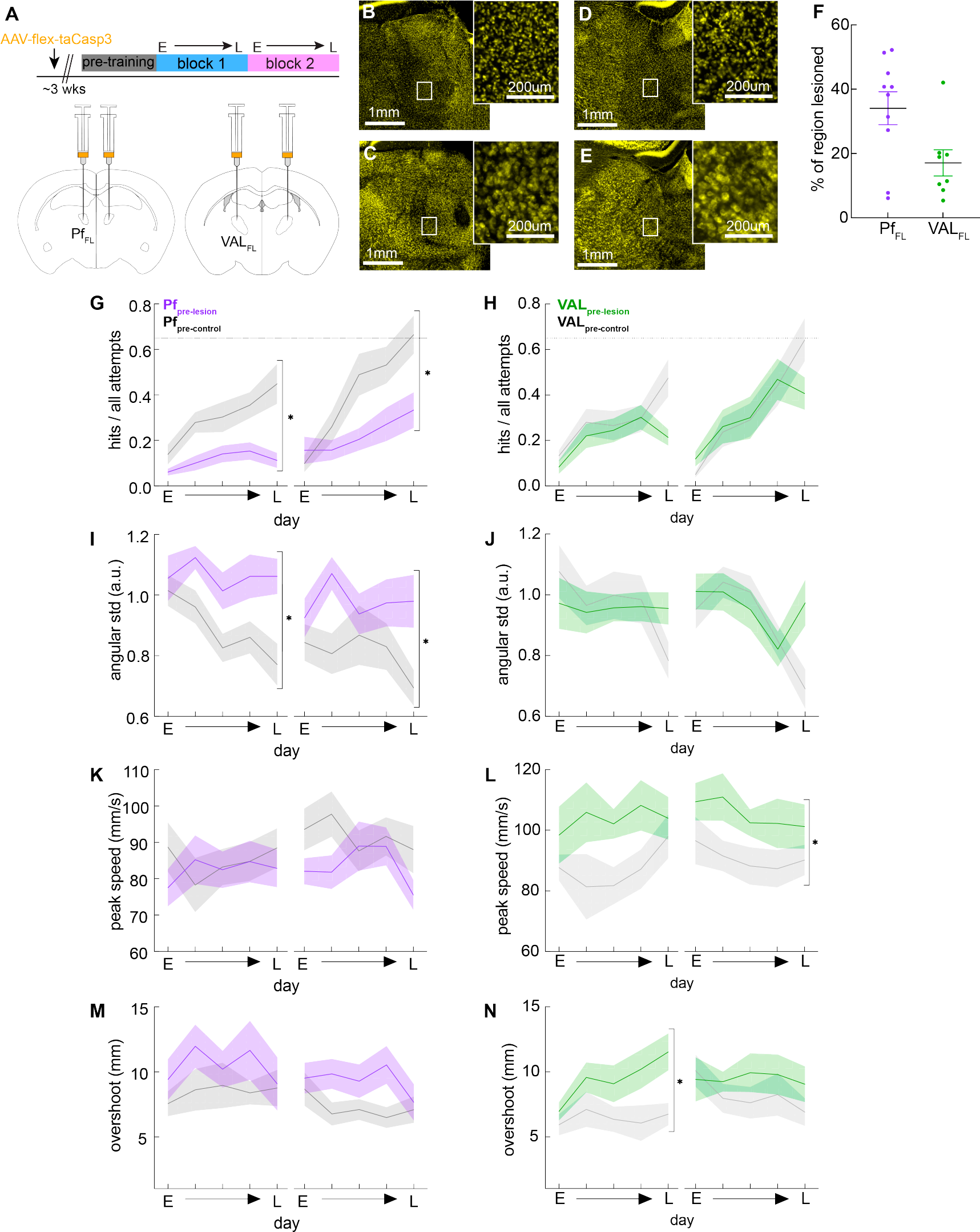
Ablation of Pf_FL_ but not VAL_FL_ cells before learning impairs the refinement of initial reach direction. (A)Experimental design showing the timeline for surgeries and STT training, and coronal ABA sections showing the injection targets for Pf_FL_ and VAL_FL_ bilateral lesions. (B) Representative coronal section of VGlut2-Cre mouse injected with AAV expressing taCasp3 -NeuroTrace (yellow) targeted to Pf_FL_.High-magnification view from the inset in box. (C) Representative coronal section of WT-littermate mouse injected with AAV expressing taCasp3 -NeuroTrace (yellow) targeted to Pf_FL_. High-magnification view from the inset in box. (D-E) Same as (B-C), for AAV injection targeting VAL_FL_. (F) Percent of Pf_FL_ and VAL_FL_ volumes lesioned, averaged for both hemispheres. Mean+/-SEM and single animals. (G) Hit ratio (hits / all attempts) on five selected days over two blocks; Early (E) day of block, Late (L) day of block, and three equidistant days in between. Pf_pre-lesion_ (purple) and Pf_pre-control_ (gray). (2-way repeated measures ANOVA: Block 1: group *p < 0.001, group × day in block *p = 0.032, day in block *p < 0.001; Block 2: group *p = 0.028, group × day in block *p < 0.001, day in block *p < 0.001. Šídák’s multiple comparison test Pf_pre-lesion_ vs Pf_pre-control_ late block 1,*p < 0.0001; late block 2, *p = 0.005) (H) Same as E, but for VAL_pre-lesion_ (green) and VAL_pre-control_ (gray). VAL_pre-lesion_ hit ratio is not significantly different than VAL_pre-control_ group. (2-way repeated measures ANOVA: Block 1: group p = 0.186; Block 2: group: p = 0.801,) (I) Variability of initial direction (angular std) for all trajectories. Pf_pre-lesion_ group has significantly higher variability of initial direction across both blocks (2-way repeated measures ANOVA: Block 1: group: *p = 0.003, day in block: *p = 0.032. Block 2: group: *p = 0.042. Šídák’s multiple comparison test Pf_pre-lesion_ vs Pf_pre-control_ late block 1,*p = 0.004; late block 2, *p = 0.032) (J) Same as in (I) for VAL lesion groups. VAL lesions do not have an effect on directional variability. (2-way repeated measures ANOVA: Block 1: group: p = 0.960. Block 2: group: p = 0.372). (K) Peak speed of all trajectories (mm/sec). Pf lesions had no effect on peak speed of movements (2-way repeated measures ANOVA: Block 1: group: p = 0.690. Block 2: group: p = 0.157). (L) Same as in (K), for VAL lesion groups. VAL_pre-lesion_ mice have higher peak speeds than VAL_pre-control_ mice across blocks (2-way repeated measures ANOVA: Block 1: group: p = 0.050, Block 2: group *p = 0.042). (M) Overshoot (mm) of target for hit trajectories. Pf lesions had no effect on target overshoot (2-way repeated measures ANOVA: Block 1: group: p = 0.216. Block 2: group: p = 0.090). (N) Same as in (M), for VAL lesion groups. VAL_pre-lesion_ mice overshoot the target more than VAL_pre-control_ animals in block 1 (2-way repeated measures ANOVA: Block 1: group: *p = 0.019. Block 2: group: p = 0.293). Šídák’s multiple comparison test VAL_pre-lesion_ vs VAL_pre-control_ late block 1: *p = 0.023.

Behavioral training began three weeks after AAV injection and all animals were trained for a maximum number of days in each block (**Figure 5A**). Pf_pre-lesion_ mice showed a strong deficit in learning to hit the rewarded target area in both blocks compared to the Pf_pre-control_ group (**Figure 5G, Figure S7A**), even though the two groups were equally engaged in the task and performed the same number of attempts (**Figure S7C**). VAL_pre-lesion_ mice, however, did not have such a learning deficit (**Figure 5H, Figure S7B**), and were also equally engaged in the task as their controls (**Figure S7C**).

These results suggest that Pf neural activity is needed to learn directional reaches.

We next asked whether the lesioned animals were able to refine the direction of movements during training. We found that Pf_pre-lesion_ mice maintained high variability in the initial direction of their reaches throughout the training, indicating that they were not able to refine their reach direction (**Figure 5I**). However, pre-learning lesions of Pf did not affect movement performance in general as speed was not affected (**Figure 5K**). In contrast, VAL_pre-lesion_ mice were able to refine their initial reach direction with learning similar to their controls (**Figure 5J**), allowing them to increase their hit ratio with training. Interestingly, we discovered that VAL_pre-lesion_ animals reached higher peak speeds (**Figure 5L**), and had larger overshoots of the target (**Figure 5N**). This is suggestive of a more cerebellar phenotype^10,19^, however, this effect on speed and targeting did not disrupt successful hitting of the target and overall performance.

These results show that both Pf and VAL lesions caused specific impairments and changes in spatial target reaches. In line with our neural decoding results, we found that lesioning Pf, but not VAL, led to a strong impairment in the refinement specifically of the initial direction of reaches, resulting in a strong learning deficit. However, lesioning VAL resulted in animals moving faster and overshooting the target more than their controls. Taken together these results reveal separable functional roles of these two thalamic nuclei in forelimb reaching movements.

### Pf and VAL are not required for the performance of learned directional reaches

Given our findings that the neural activity and direction prediction accuracy was strongest during early days in learning, but decreased or disappeared on late days with more refined movements, we also hypothesized that these thalamic areas are required early in learning but not for the execution of already learned reaches. To this end, we again ablated glutamatergic cells in Pf_FL_ and VAL_FL_ in Vglut2-Cre mice and used their WT littermates as controls, but only once they had learned to reach to the first target. First, mice were implanted with a headcap and headpost and trained in block 1 of the STT for 28 consecutive days. After block 1 training, all mice were bilaterally injected with AAV5-flex-taCasp3-TEVp (AAV5) in either Pf_FL_ or VAL_FL_ (Pf_post-control_ n = 5, Pf_post-lesion_ n = 6, VAL_post-control_, n = 6; VAL_post-lesion_ n = 4) (**Figure S8A-C**). After a three-week break to ensure full expression of Caspase3, mice were trained for an additional nine test days on the same target from block 1 (**Figure 6A**).

**Figure 6.**
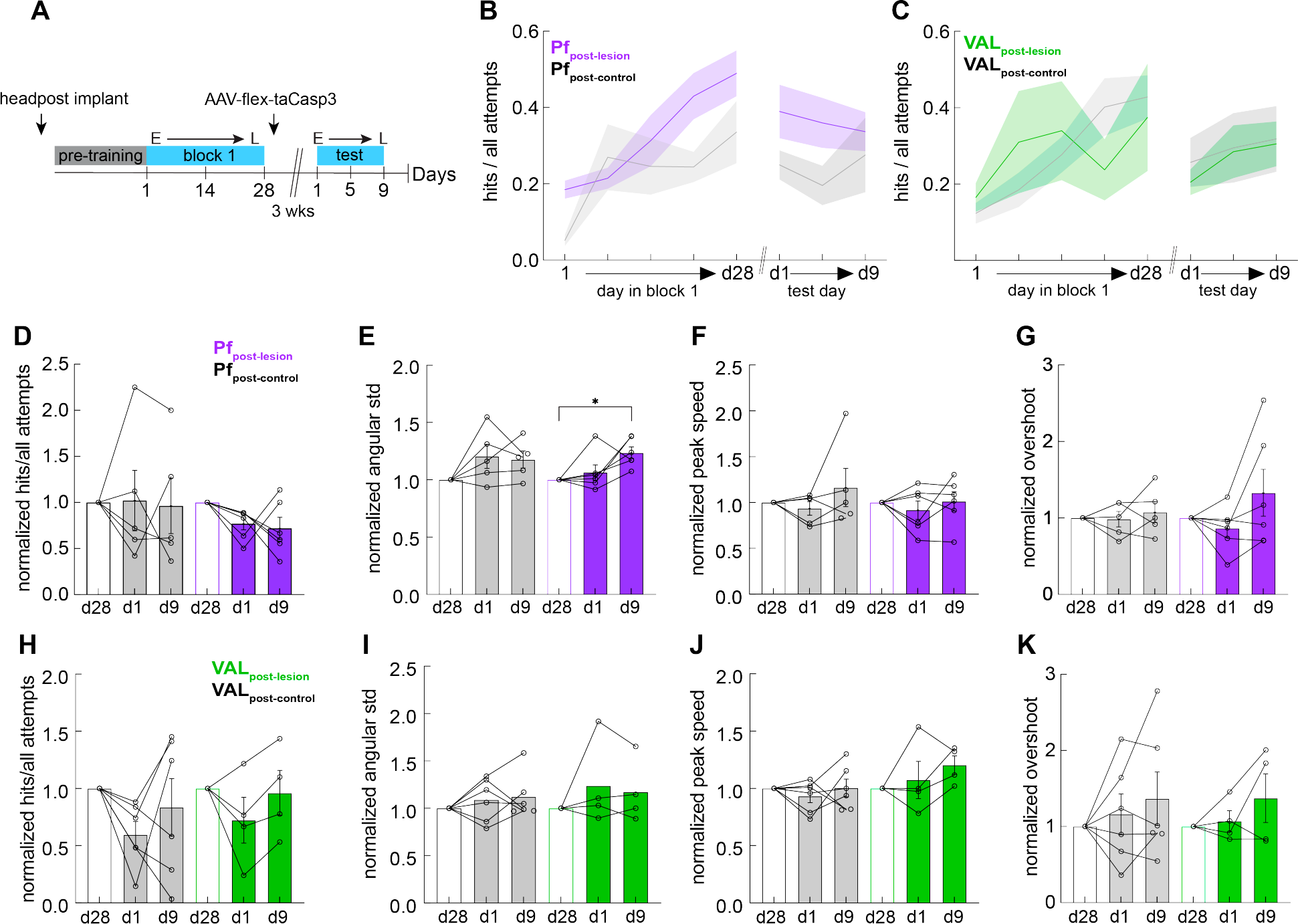
Pf and VAL are not required to perform well-learned directional reaches. (A) Experimental design showing the timeline of surgeries and STT training and testing. (B) Hit ratio (hits / all attempts) on five selected days for block 1 and three test days post-lesion. Pf_post-lesion_ (purple), n = 6 mice; Pf_post-control_ (gray), n = 5 mice. Mean ± SEM in thick colored lines and shaded bounds. (C) Same as in (B), for VAL lesion groups. VAL_post-lesion_ (green), n = 4 mice; VAL_post-control_ (gray), n = 6 mice. (D) For Pf_post-lesion_ (purple) and Pf_post-control_ (gray) groups, normalized performance on the last day of block 1 before lesions (d28), and the first (d1) and last (d9) day of post-lesion test. (Wilcoxon test for d9 vs d28: Pf_post-lesion_, p = 0.125; Pf_post-control_, p = 0.813) (E) For Pf_post-lesion_ (purple) and Pf_post-control_ (gray) groups, normalized variability of initial direction. Pf_post-lesion_ mice increased their directional variability after lesions (Wilcoxon test for d9 vs d28: Pf_post-lesion_, *p = 0.031; Pf_post-control_, p > 0.999) (F) For Pf_post-lesion_ (purple) and Pf_post-control_ (gray) groups, normalized peak speed of movements. There is no effect of Pf lesions. (Wilcoxon test for d9 vs d28: Pf_post-lesion_, p = 0.750; Pf_post-control_, p = 0.813) (G) For Pf_post-lesion_ (purple) and Pf_post-control_ (gray) groups, normalized overshoot of target. There is no effect of Pf lesions. (Wilcoxon test for d9 vs d28: Pf_post-lesion_, p = 0.688; Pf_post-control_, p = 0.813) (H-K) same as in D-G, but for VAL_post-lesion_ (green) and VAL_post-control_ (gray) groups. There is no effect on any metric in the VAL_post-lesion_ group. (for VAL_post-lesion_ Wilcoxon test d9 vs d28, p = 0.875 for normalized hit ratio; p = 0.500 for normalized variability of initial direction; p = 0.125 for normalized peak speed; p = 0.625 for normalized overshoot) (D-K) performance metrics on the first (d1) and last (d9) day of post-lesion test. Data normalized to the values on the last day of training before lesion (d28, open white bars). Filled colored bars denote post-lesion days for Pf_post-lesion_ (purple bars), VAL_post-lesion_ (green bars), and control groups (gray bars) Individual animal data plotted with open circles. Mean ± SEM shown.

To examine how performance of directional reaches was affected by lesions after learning (**Figure 6B**), we normalized all test day behavior to the last day of pre-lesion training and compared the animals performance to its baseline. This analysis revealed that the thalamic lesions did not affect the successful performance of directional reaches (hit ratio) (**Figure 6D/H**), nor did they affect the engagement of the animals in the task directly after the lesion (number of attempts made) (**Figure S8D/F**). Interestingly, Pf_post-lesion_ mice significantly increased the variability of their initial reach direction compared to baseline at the end of the 9 day testing period (**Figure 6E**), as well as increased the number of attempts made in a session (**Figure S8D**). These effects were not seen in the control group or the VAL cohort (**Figure 6I**). This may indicate that the lack of Pf activity gradually impaired the precision of the initial direction of reaches over many days of training. Still, we did not detect such a trend for the peak speed of movements or the target overshoot in the VAL animals (**Figure 6J/K**), nor the Pf animals (**Figure 6F/G**). These results suggest that post-learning VAL is not needed to regulate reach speed once target reaches have been learned. However, post-learning Pf lesions led to long-term effects in the precision of learned directional reaches, suggesting that Pf activity may be needed to maintain low directional variability with continued training over long periods of time.

## DISCUSSION

Through functional recording and manipulation experiments, we revealed that Pf_FL_ and VAL_FL_ have dissociable roles in the refinement and execution of forelimb reaches to a spatial target. We mapped Pf_FL_ and VAL_FL_ as the thalamic areas that primarily target forelimb striatum and the CFA in M1, respectively. Pf_FL_ and VAL_FL_ neural populations were most active during reaching on early days, but the magnitude of their engagement decreased over learning, which was mediated through a change in the number of reach-responsive cells. This result was further confirmed when tracking the same cellular populations over multiple days, where the matched cells increased their modulation after a target switch. We also found, using PCA, that the dimensions capturing the dominant co-activation patterns of the same matched neural populations in Pf_FL_ shifted between early and late stages of learning, but stabilized after learning. In contrast VAL_FL_ dynamics shifted from day to day regardless of learning stage. Importantly Pf_FL_, but less so VAL neurons, encoded directional information early in learning, but that directional information was not present late in learning. In line with these findings, targeted pre-learning lesions of Pf_FL_ and VAL_FL_ revealed that Pf_FL,_ but not VAL_FL_, is needed to refine directional variability of movements and consequentially to learn to reaches the target. VAL_FL_ lesions, on the other hand, did not affect the learning and refinement of the reaches but affected the speed of movement and led to animals overshooting the target. Overall, this data is aligned with previous findings showing that Pf and VAL have important roles in motor behavior^3,21,44,45^, but reveals an exquisite dissociation in their roles in learning to refine directional reaches to a target.

General movement related activity in Pf was seen in mice during performance of behaviors such as lever pressing, rotarod, and reward-tracking assays^25,44,46^. Our work complements and expands on these findings, which focus on activity during movements already well-learned, by closely examining how Pf activity evolves over the learning process and what behaviorally relevant information can be decoded by neural populations. Other work has shown that VAL projections to M1 dynamically reorganize while training grasping movements^47^, and are needed to perform cued grasping and lever-pulling tasks after learning^3,45^. With our focus on learning over time, we’ve added to these findings that VAL_FL_’s low-dimensional activity changed regardless of learning stage, with individual cells changing their modulation from day-to-day. Our neural analysis revealed that that Pf and VAL undergo significant changes with behavioral refinement and learning of directional reaches, and are preferentially engaged in early learning.

One of our most striking results came from investigating if Pf or VAL neural activity encoded the direction of reaches. The thalamus is already known to contain head direction cells, which encode animal’s directional heading in the horizontal plane. Head direction cells are found in the anterior dorsal thalamic nucleus, as well as in other areas distributed across the brain like the lateral mammillary nuclei, entorhinal cortex, and dorsal striatum^48–51^. Recent work reported that Pf has a role in controlling the direction of head turning actions in a reward tracking task in mice^46^. Our work in the forelimb domain aligns with these findings, and expands on them to include Pf in regulating direction of reaching actions. Interestingly, Pf activity best encoded directional information on early learning days, and could not be used to decode direction late in training after learning occurred. The upstream source of this directional information may stem from the inputs to Pf from the superior colliculus, which is thought to send an orienting signal^52,53^. Unpredicted or especially significant events cause short-latency responses in the superior colliculus and Pf, which can be quickly routed to the striatum by way of Pf ^54^. These signals may potentially serve as general directional information used to guide behavior.

Recent work in mice has highlighted Pf as having a specific role in learning^21,25,55^. Our findings are overall consistent with these previous reports, but gives more insight into what behaviorally relevant information Pf contributes during learning, which we found to include the direction of forelimb reaches. Consistent with this directional information in Pf_FL_ neural populations, our lesion studies support our hypothesis that Pf activity is critical to refine the direction of the target reaches. When we performed pre-learning lesions of Pf_FL_, mice did not reduce the directional variability of their movements, and as such did not refine reaches to targets. However, these pre-learning lesions did not affect the variability of directional reaches early in training, indicating that Pf is not needed for the exploration of different directions. Rather, our findings support a function of Pf in conveying directional information to downstream structures which reinforce behavior, such as the striatum. An interesting hypothesis would be that Pf sends information about the direction of forelimb movements to the striatum early in learning, allowing for successful direction commands to be reinforced through dopamine release upon reward delivery. Later in training after refinement, when the reinforced direction has been learned and directional variability is low, Pf conveys less directional information. Interestingly, when we performed post-learning lesions of Pf_FL_, mice were still able to perform with the same hit ratio, but after nine days of training post-lesion, the directional variability increased. This suggests that Pf activity is required for the maintenance, and continuing reinforcement, of the appropriate direction of forelimb reaches. These ideas align with recent findings that Pf inputs to the striatum are needed to learn and maintain the performance of a sequential level pressing task^21^.

In contrast to Pf_FL_, we found that VAL_FL_ activity did not encode direction as strongly, nor was it needed to learn or perform targeted reaches. However, as our lesions left areas of VAL intact, it is possible we would see different behavioral effects with more of VAL lesioned. Previous work detailed that VAL input to M1 was needed for mice to learn a lever pulling task^45^. The differences in our findings may be explained by task requirements, as their behavior required head-fixed mice to pull and hold a lever for a set amount of time. In our task, success was not contingent on the speed of movements and mice were not required to stop their movements inside of the target. As such we found that mice with pre-learning lesioning of VAL_FL_ moved faster and overshot the target more, but they were able to increase their hit ratio by learning the direction to the target. There was also a subtle, but not statistically significant difference, in the hit ratio of VAL lesioned mice at the late stage of training, which may hint that the final refinement needed for higher precision did not occur. These results suggest that VAL may play a role in more sensory error based learning and online correction of actions^56,57^. VAL lesioned mice, with an intact Pf, learned the direction of movement, which was enough to increase the number of rewarded actions. Future experiments will explore whether VAL lesioned mice could learn a reaching movement that required more endpoint precision. This is an interesting line of future work, as it aligns well with documented deficits in endpoint precision in human patients with cerebellar damage^19^, as well as recent work in mice linking cerebellum to endpoint precision in reaching tasks^9,10,58^.

Our findings support the overarching hypotheses that the basal ganglia is involved in motor learning through reinforcement learning, whereas the cerebellum is more important for movement initiation and error-based learning^59–61^, and open the door for future studies to address the relative roles of sub-populations of Pf or VAL. Furthermore, recent findings demonstrated how subpopulations of Pf have specific behavioral functions, showing that Pf projections to other areas besides the striatum, like the subthalamic nucleus, have different roles in motor behaviors^25^. These subpopulation-specific functional roles are important to consider to gain a comprehensive understanding of system-wide models of behavior. Additional investigation using projection specific recordings or manipulations in our task may reveal even more detailed distinctions of Pf and VAL roles in reaching movements.

Taken together, our work shows that within the thalamus, there are multiple circuits governing separable components of motor behavior and refinement, namely direction and speed, with learning. Not only are Pf and VAL performing different functions, but these roles dynamically evolve with learning, which has important implications for understanding motor learning and also movement related disorders.

## Lead Author

Further information and requests for material and codes should be directed to and will be fulfilled by the Lead Contact, Rui M. Costa

## Data code and availability

All analysis were performed using publicly available software and custom scripts written in MATLAB R2019b (Mathworks) and Datajoint (Datajoint 0.13.0). The data of this study and codes are available from the authors upon reasonable request.

## MATERIALS AND METHODS

### Mice

All experiments and procedures were performed according to National Institutes of Health (NIH) guidelines, with lab protocols approved by the Institutional Animal Care and Use Committee of Columbia University. Adult male mice (n=10), aged 3-6 months, were used for all 2-photon calcium imaging experiments. Adult mice of both sexes (n=25 female, n=32 male), aged 2-6 months, were used for all lesion experiments. Adult mice of both sexes (n=3), aged 4-8 months were used for anatomy experiments. The strains used were: C57BL6/J (Jackson Laboratories, strain #: 000664), VGlut2-Cre (Jackson Laboratories, strain #: 028863, C57BL6/J background). Prior to surgery, adult mice were housed in groups (2-5 per cage). Following surgery, mice used for behavioral experiments were individually housed with running wheels for enrichment. All mice were kept under a 12-h light–dark cycle.

## METHOD DETAILS

### General surgical procedures

Analgesia was administered in the form of subcutaneous injection of Buprenorphine SR (0.5-1 mg/kg) on the day of surgery. Mice were anesthetized with isoflurane (1%-5%, plus oxygen at 1-1.5 l/min) and then placed in a stereotaxic holder (Kopf Instruments). The scalp was shaved and cleaned with 70% alcohol and iodine, and intradermal Bupivacaine (2 mg/kg) was administered. The mouse’s body temperature was maintained throughout surgery at 37°C using an animal temperature controller (ATC2000, World Precision Instruments). After surgery mice were allowed to recover until ambulatory in the home cage on a heating pad.

### Viral injections

For **anatomical tracing** experiments, a midline incision was made to expose the skull, and a craniotomy was made over the injection sites. Cholera toxin subunit B (Invitrogen lot #2384712, Alexa Fluor 555 conjugate; Invitrogen lot #2387465, Alexa Fluor 488 conjugate) reconstituted at 1% in HEPES buffered saline. Injections were made at the following volumes in respective areas (coordinates in mm from bregma): DLS at 0.75 AP, 2.45-55 ML, 2.45-5 DV: 80-100 nl; CFA at 1.5 AP, 2.3 ML, 0.55 DV: 45 nl.

**Imaging animals** were unilaterally injected with 200-300 nl AAV5-CamKII-GCaMP6f-WPRE.SV40 (UPENN — lot #CS1250, titer: 4.12x10^12^ GC/mL; or Addgene — lots #v23807 and #v59618, titer: 2.3x10^13^ GC/mL), or AAV5-CAG-flex-GCaMP6f-WPRE.SV40 (Addgene — lot #v28551, titer: 3.5x10^13^ GC/mL) in Pf or VAL (Pf at -2.0 AP, 0.88 ML, 3.6 DV; VAL at -0.86 AP, 0.9 ML, 3.6 DV). A subset of animals were also injected with 60-100nl retro-tdtomato virus pAAV-CAG-tdTomato (Retro) (ZIVirology custom virus— titer: 8.1x10^12^ vg/mL, lot #8) into either the forelimb-related area of DLS or the CFA. The expression of tdtomato was used for additional post-hoc verification of targeting forelimb projecting regions.

**Pre- and post-learning lesion** animals were bilaterally injected with 35-50 nl of AAV5-Ef1a-Flex-taCasp3-TEVp (UNC – lot #AV5760G, titer: 4.2x10^12^ vg/mL) in Pf or VAL at the same coordinates reported for imaging animals. All viruses and tracers were injected with the Nanoject III Injector (Drummond Scientific, USA).

Headpost and tube rim implantation

All behavior animals followed the same pre and post-surgery protocol as described above. After mice were anesthetized and placed in a stereotaxic holder the scalp was removed to expose the skull. The fascia was cleared off and the cranium cleaned with saline and 3% hydrogen peroxide. A custom metal headpost (ZI Advanced Instrumentation platform) was secured to the skull using C&B Metabond dental cement (Parkell). The headbar was designed with small U-shaped ends on either side of the straight cemented bar that allowed easy sliding in and securing of the mouse’s head in the head-fixation holders by tightening a screw through the U-shaped ends.

For post-learning lesion animals, mice underwent two surgeries. For surgery one, mice were implanted with a sterilized PCR tube rim (Axygen PCR strip tubes; REF: PCR-0212-FCP-C) and headpost. The tube rim was used to create a reference coordinate system to level the brain in surgery two, and to keep the skull above the injection target areas free from dental cement. The rim was placed centered around bregma, and secured with a combination of Metabond dental cement and superglue (Loctite). After the rim was secured, the headpost was implanted. Measurements were taken between the rim’s most anterior and posterior points aligned with bregma, two points of the headpost, and the marked anterior-posterior and medial-lateral coordinates for Pf or VAL. The exposed skull was again thoroughly sterilized, and then covered with a non-toxic sealant (Kwik-Sil silicone sealant). Surgery two took place after initial STT training. Because lambda was no longer visible after surgery one, the new coordinates measured at the end of the first surgery were used to level the mouse’s skull in the stereotaxic frame. Craniotomies and viral injections were then made in the same manner as described above. Surgery two was completed by filling in the leveling rim with dental cement.

### Chronic GRIN lens implantation

After viral injection as described above, a 0.5 mm diameter gradient index (GRIN) lens (length: 6.51 mm, working distance: image side: 0 μm in air, object side: 500 μm in water, design wavelength: 920 nm, NA 0.5, non-coated, NEM-050-50-00-920-S-1.5p, GRINTECH) was implanted in the left thalamus (contralateral to the arm moving the joystick) above the injection site. Once in place, the lens was secured to the skull using a combination of Metabond dental cement and superglue. After GRIN lens implantation, headposts were implanted as described above.

### Behavioral setup

SCARA joystick hardware and Spatial Target Task controls The SCARA (Selective Compliance Articulated Robot Arm) joystick and rig was built as described previously^33^. Animals were head-fixed in a custom-designed 3D printed cup (copyright IR CU21353), that we have previously shown allows increased workspace exploration compared to the standard tube. All animals used their right forelimb to interact with the SCARA joystick. A metal screw was placed horizontally to the left of the joystick to be used as arm rest for the left limb. The Spatial Target Task (STT) was controlled through a microcontroller board (Teensy 3.6, Arduino) and breakout board platform custom-built by the Advanced Instrumentation Platform at the Zuckerman Institute (TeenScience). The behavior task was written using the Arduino IDE. Briefly, the joystick was actively moved, or kept at the start position through a proportional integral derivative (PID) algorithm. The joystick position and all task events were recorded via serial output commands and recorded through Bonsai (OpenEphys).

### Spatial Target Task design and training

Animals were food restricted and given an individualized amount of chow food after each training session to maintain their body weight at 80% of pre-training baseline. Each session lasted until a maximum number of rewards (5-7 μl drop of 7.5% sucrose in water) were achieved, or the maximum time had elapsed.

During the session, the animal could move the joystick out of the start position (0/65 mm from the motor axis, 1 mm radius) and explore the workspace without any force generated by the motors for 7.5 sec per attempt in a self-paced uncued manner.

The training schedule was designed as described previously^33^ and included 2 pre-training phases, and 2 blocks of target training. Briefly, pre-training consisted of 4 days of Phase 1, and 5 or more days of Phase 2. In Phase 1 of pre-training, each initial touching of the joystick was rewarded (with a delay of 500 msec on days 1 and 2 and 1000 msec on days 3 and 4). Rewards were also given at random intervals between 5 to 15 sec for continuous touching of the joystick, until 100 rewards were achieved.

For both Phase 2 and the target training, animals needed to move the joystick out of the start position and explore the workspace to receive a reward. If the mouse let go of the joystick for > 200 msec, the attempt ended, the motors engaged and moved the joystick back to start (miss). If the criteria for a reward was met, the reward was delivered through the solenoid and the motors engaged and moved the joystick back to the start after a 750 msec delay (hit). In Phase 2 a reward was given for moving the joystick in a forward direction of either a 40° or a 60° segment. Using the rewarded forward trajectories on the last day of Phase 2, 2 target locations were defined for each animal (**Figure S1B**). The mean initial direction of all rewarded trajectories was calculated and a target center defined 40° to the left and right of the mean direction at 7.5 mm distance from the start position with a target radius of 3 mm for imaging animals, and 8mm distance from the start position with target radius of 2.75 mm for lesion animals (**Figure S1A**).

#### Target training

Target 1 was rewarded during the first block, and target 2 during the second block of target training. The reward was delivered immediately upon entering the target circle. A short ITI period, enforced by the SCARA motors, required the animal to stay in the start position between trials. The ITI could only be exited by exerting less than 11 g force against the joystick in any direction. After the ITI, each movement of the joystick out of the start position by the mouse was counted as an attempt. Task performance was calculated as the number of hits divided by all attempts (hit ratio). For imaging studies, when animals reached a 3-day average hit ratio > 0.6 or reached a maximum number of training days (53 days) on block 1, the target was changed in the next session and a new block began (performance criterion). For pre-learning lesion studies, maximum number of training days were 35 days in block 1 and 19 days in block 2. For post-learning lesion studies, mice were trained for consecutive days before lesions and 9 additional days post-lesion on the same target.

#### Behavioral training during 2-photon calcium imaging

To optimize head orientation under the 2-photon microscope, each mouse was trained with its own head-fixation setup, specifically adjusted for its headpost implantation. Pre-training took place in sound attenuating behavioral boxes for all mice. Mice were moved into the 2-photon behavioral rig after reaching over 0.5-0.65 hit ratio in pre-training. For both Pf and VAL cohorts, 2 mice each were moved into the 2-photon only after starting block 1. Performance often dropped once mice were moved into the 2-photon behavioral setup. Daily training sessions under the 2-photon microscope lasted until 50 or 120-150 rewards (on early and late days respectively) were achieved, or a maximum time had elapsed (60-80 min). For low performing animals, the target radius was increased early in block 1 or block 2, to up to 4.5 mm, but was reduced back to 3 mm by the end of the block.

### Analysis of Spatial Target Task behavior

All data was analyzed using custom MATLAB code (MATLAB engine for python R2019b, Mathworks, Inc.) run from a Python analysis pipeline (Python 3.7.8) through a custom DataJoint^62^ database (Datajoint 0.13.0). Data from the target training blocks are shown using 5 selected days per block. For each block that includes an early and a late day and 3 equidistant days in between. Two-way ANOVA with repeated measures with Šídák’s correction, one-way ANOVA with repeated measures with Šídák’s correction, paired and unpaired t-tests, Chi-squared test, and simple linear regression were used for statistical analysis. A p-value of less than 0.05 was considered statistically significant.

#### Preprocessing of trajectories

Trajectories were down sampled to 6 msec intervals.

*Trajectories used for quantification:* For each attempt a joystick trajectory was recorded. All trajectories were used to quantify refinement of the movement. For hit trajectories, only the path from start to the point of target entry was used in the analysis of the average trajectory variance.

### Calculation of trajectory metrics

Behavior metrics were generated from raw joystick trajectories and analyzed as previously described^33^.

Briefly, the *Occupancy* of the workspace was quantified across 1 x 1 mm bins using the MATLAB ‘histcount2’ function on each trajectory. Dwell time in each bin and multiple visits to the same bin were discounted to calculate the total area visited, but are shown as heatmaps for an example animal (Figure 1G). To calculate the *mean trajectory variability* trajectories were downsampled to 200 points. The mean trajectory was calculated by averaging the 200 coordinates of all trajectories. Then the standard deviation was calculated as the square root of the squared shortest distance to the mean trajectory at each point, divided by the number of trajectories. The average standard deviation along the 200 is reported.

The *initial vector variability* was calculated from initial vectors defined from the point at which the trajectory left the start circle (1 mm radius) to the point of it crossing a circle of 2.2 mm radius from the start position center. For each vector the angle was calculated using the ‘atan2’ MATLAB function. The angular standard deviation across all angles was calculated using the ‘circ_std’ function from the CircStat circular statistics toolbox^34^, which is bounded between the interval *[0, √ 2]*. The *peak speed* was calculated on the downsampled (6ms) trajectory, which was smoothed using a 30 msec moving average (‘smooth’ function in MATLAB). For analyzed hits, only the trajectory from the start until entering the target was considered. The maximum value of the smoothed speed of each trajectory was averaged across all trajectories.

To calculate the *target overshoot* the pathlength between the point of the trajectory entering the target and the end of the trajectory, when motors engaged (750 msec after entry), was measured. The average target overshoot pathlength per session is reported.

### Two-photon calcium imaging

All imaging experiments were conducted on a Bergamo II rotatable two-photon laser-scanning microscope (Thorlabs, Inc.). The system was configured with 8 kHz resonant-galvo-galvo laser scanning mirrors and imaging frames of 512 x 512 pixels (corresponding to an area of 834 μm x 834 μm) were acquired at 30 fps. The system was equipped with two-channel fluorescence detection with amplified non-cooled GAsP photomultiplier tubes (PMTs). Emitted fluorescence was first directed to the PMTs, then split into “green” and “red” channels by a 565 nm sharp edge long-pass dichroic mirror. The green channel and red channel were subsequently filtered by a 525 nm/39 nm bandpass filter, and 593 nm /40 nm bandpass filter, respectively, before detection in the PMTs. The microscope was controlled via ThorImage 4.

Imaging was performed through a Nikon 16x, 0.8 NA water immersion objective placed over the implanted GRIN lens with the laser beam sized to fill the back aperture. A Coherent Chameleon Vision S tunable titanium-sapphire laser tuned to 920 nm with 75 fs pulses at 80 MHz repetition rate was used. Dispersion correction was adjusted to maximize fluorescent brightness as recorded under the objective. The imaging power was modulated through a Conoptics 350-105 Pockels Cell driven by a Conoptics 302 RM Amplifier for each imaging session to stay in the linear range of pixel intensity. The same field of view was imaged across days of behavioral training by matching the depth of imaging across sessions. To identify imaging fields of view, head-fixed mice were positioned such that their GRIN lenses were aligned to the imaging objective. The depth at which GCaMP fluorescence signals were strongest was noted for each individual animal, and each following imaging session was done at that approximate identified depth.

### Synchronization of behavioral and imaging data stream

Behavioral and imaging data were synchronized in two ways. Behavior events (attempt start, reward delivery, joystick touch) were recorded as TTL pulses together with the imaging frames through a National Instruments DAQ and saved to an h5 file using ThorSync (Thorlabs, Inc.). The h5 file datastream wasthen used to synchronize with the full behavioral data using custom MATLAB code. Further, a TTL pulse (∼10 μs) from the microscope was sent directly to the TeenScience behavior acquisition board for each imaging frame, directly integrating the imaging frame timestamp into the behavior datastream.

### Calcium imaging data pre-processing

Calcium imaging data was motion corrected using non-rigid motion correction, cells were segmented and raw fluorescence traces extracted using Suite2p (0.10.0)^42^. Automatically classified cells were further manually curated using the Suite2p GUI and cells with a skew of less than 0.1 were excluded. Using custom MATLAB code, the neuropil signal was subtracted from the raw fluorescent traces using a coefficient of 0.7. The trace was then detrended by subtracting the baseline fluorescence. Baseline fluorescence was calculated on the full session’s trace using a gaussian filter and minimum followed by maximum filtering. After detrending, individual neuron traces were z-scored across the entire imaging session. This z-scored trace was used for all following analyses.

### Population activity analysis

To compute peri-event trial-averaged activity, the neural activity of individual neurons was aligned to the event and then averaged across attempts. Three different event windows were defined: pre-movement (-500 – 0 ms from movement onset), during movement (0 – 500 ms from movement onset), and after hit (0 – 500 ms from target hit). If the trial-averaged trace for a cell was above or below 3 x SD of that cell’s baseline trace (trial-average from -750 – -600 ms before movement onset) for over 90 ms, that cell was classified as significantly up- or down-modulated, respectively. Each cell was classified as significantly modulated or not during each of the three time windows (pre-movement, during movement, and after hit). To compare the average population activity during the movement across sessions of the block, the trial-averaged activity of all cells from all animals of the Pf or VAL groups were pooled and averaged.

### Tracking cells across multiple imaging sessions

Cell matching was performed across two pairs of sessions; a late day of block 1 to the early day of block 2, and across two late days in block 1. We imaged the same field of view across experimental sessions based on landmarks in the field of view, and developed a method to identify which cells in a chosen session were present in the other session. Cell matching was performed using the motion-corrected, pre-processed, and curated region of interest (ROI) masks from the Suite2p analysis. One session was chosen as the “template session”, and the ROIs of the other session were aligned to this template session. Specifically, we used Suite2p’s non-rigid motion correction method to align the average image of each session to the average image of the template session, and applied the corresponding alignment shifts to the ROIs identified in the session. Then for all sessions, we calculated the percent-overlap between each ROI in the template session and the most overlapping ROI in the other session. We used each ROI’s binary mask, i.e., the image of the ROI where each pixel had the value 1 if it was part of the ROI and the value 0 if not, and we calculated the percent-overlap between two ROIs (ROI A and ROI B) as the average of two quantities: 1) the percentage of ROI A’s pixels that are also ROI B’s pixels, and 2) the percentage of ROI B’s pixels that are also ROI A’s pixels. We determined that an ROI on one session was matched to an ROI on another session if the percent-overlap exceeded 55%. This 55% threshold was based on our visual inspection of example sessions across animals (**Figure S4**). Thus, this procedure determined if each ROI from a template session matched to a ROI from another session. Finally, we applied this procedure to calculate which ROIs on the template session were present on the other chosen session and used only those cells for analysis. Cell matching resulted in the following numbers of matched cells: late block 1 vs. early block 2: Pf: n = 116 cells, 4 mice, VAL: n = 86 cells, 5 mice; mid-late block 1 vs. late block 1: Pf: n = 108 cells, 4 mice, VAL: n = 72 cells, 4 mice.

### Principal component analysis for matched cells

For visualization of neural activity (**Figure 3D-G**), principal component analysis (PCA) was performed on matched cells from the late day of block 1 (L) and early day of block 2 (E) combined for all animals of the group, and in a second condition comparing matched cells from a mid-late day of block 1 (ML) and late day of block 1 (L). PCA was used to calculate the dimensions in which neural population activity most varied within neural activity space. In pre-processing, each cells’s fluorescence was de-trended (see Methods) and z-scored based on the fluorescence activity across the entire session. We extracted movement-locked neural activity as the trial-averaged activity, i.e., the peri-event time histogram or PETH, in the temporal window starting 2 seconds before movement onset and ending 2 seconds after movement onset, creating a N_frame_ x N_cell_ matrix per day. For the visualizations in **Figure 3D-H** and **Figure S5D-G**, the matrices from the days with matched cells were vertically concatenated to build the full matrix *X*. PCA was performed on matrix *X* using the MATLAB function ‘pca’. The returned scores of the first 3 PCs were plotted separately for the first half of the frames (corresponding to day L1 of ML1) and the second half of the frames (corresponding to day E2 or L1).

We then assessed whether the dimensions of neural variance were more different across two conditions than expected within a condition, separately analyzing each animal’s simultaneously recorded neural population. We subsampled trials so that each condition had the same number of trials (i.e., the number of trials in the condition with fewer trials), and we randomly divided each condition into two halves of trials. For each condition-half, we calculated the movement-locked PETH, the covariance matrix |Σ between the |*U* neurons across time based on the PETH, and performed PCA on the covariance matrix to identify the dimensions that capture the most variance. We extracted the covariance within the 3 dimensions capturing the most variance, i.e., within the subspace spanned by the top 3 principal components (PCs). To do this, we calculated the singular value decomposition (SVD) of the covariance matrix *U* ∈ *R*^*N×N*|^ where *U* ∈ *R*^*N×N*|^ contains the PCs as column vectors and |*U*_1:3_ ∈ *R*^*N*^| is a diagonal matrix containing the singular values which are the variance within each PC. The covariance within the top 3 PC’s was calculated as |*U*_1:3_|, ∈ *R*^*N*^ where *U*_1:3_ ∈ *R*^*N*^ is the matrix containing the top 3 PCs as column vectors and |*B*| is the diagonal matrix containing the singular values corresponding to the top 3 PC’s. We then calculated an alignment index of this low-dimensional covariance between the two condition-halves within a condition and compared it to the alignment between condition-halves across conditions.

The alignment index was the fraction of low-dimensional variance in condition-half |*B* captured in the subspace of condition-half *B* We projected the low-dimensional covariance of condition-half |*P*_*B*_ into the subspace of condition-half |*P*_*B*_ as |*P*_*B*_, =*U*_1:_, where 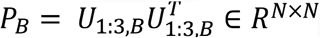 is the projection matrix into the subspace spanned by condition-half B’s top 3 PC’s. We then calculated the fraction of variance captured in the subspace as: |*B* Our alignment index took the average of the alignment of |***B***| with |*B* and the alignment of |***A*** with, as these two calculations are not equal in general. (For intuition of this asymmetry, we consider an example with two dimensions. Imagine PC1 is the same for |*A* and |*B* but that PC2 is orthogonal for *B*| and *B*, and that the fraction of |***B*** variance in PC1 is more than |*B*. In this example, the fraction of low-dimensional variance |***B*** in subspace |***B*** is higher than the fraction of low-dimensional variance |***A*** in subspace ***A***|, because |***A*** concentrates more of its variance in the dimension that is shared across |***B*** and |***B***.) We calculated the alignment index between condition-halves within a condition (and then averaging over the two conditions) and across conditions (averaging over the two directions of alignment) for each animal, averaging over 100 folds of matching trials across conditions and dividing conditions into halves.

In addition, we calculated a shuffle alignment index (for both “within-condition” and “across-condition”) to assess a chance rate of alignment. In order to preserve the temporal structure of population activity but randomize the subspace in which temporal activity evolved, we shuffled the neuron identity of the |***B*** covariance matrix before calculating the PCA and performing the within-condition and across-condition alignments: |× *β*| This shuffle alignment allowed us to assess whether alignments in the data were greater than expected by chance.

### Decoding of movement initial direction

Ridge regression models (sklearn.linear_model.Ridge, scikit learn) were used to predict the direction of initial vectors of movement (see Methods: *initial vector*). Neural data from the first 166.66 ms (5 frames) following movement initiation of all cells was concatenated (X = N_trials |×_(N_roi_ *5 frames)) and used to predict the sine and cosine of the initial direction angle for each trajectory (Y = X ×*β*). To correct for differences in variability in the dependent variable (initial direction), we subsampled the trials of all sessions to match the lowest variability of initial angle (angular std) on the late day of block 1. One session was excluded because this subsampling left less than 50 trials for model training. Data was split in 70% train and 30% held-out test trials and 8-fold cross-validation on the train dataset was performed for hyperparameter model selection. For each session 10 models were trained on the train dataset of 10 different train/test splits and used to predict the sine and cosine of the initial angle of the held-out test trials. Model accuracy was calculated using the coefficient of determination (R^2^) between the true initial direction and the model prediction. R^2^ values were averaged across the 10 models for each session.

### Immunohistochemistry

Mice were deeply anesthetized with isoflurane and transcardially perfused with 1x phosphate-buffered saline (PBS) followed by ice-cold 4% paraformaldehyde (PFA). Brains were extracted and post-fixed in 4% PFA overnight, and then transferred to PBS. Brains were sectioned into 50-75μm coronal sections using a vibratome (Leica vibratome VT1000). Sections were either stored free-floating in PBS at 4°C, or serially mounted onto slides, dehydrated and stored at 4°C until they were processed. All sections were first rinsed with PBS and permeabilized with 1-3% Triton in PBS. Immunostaining for GCamp on slide mounted sections was performed with primary antibodies (Anti-GFP Polyclonal Antibody, Alexa Fluor 488; ThermoFisher Scientific) diluted at 1:500 – 1:1000 for 1 day at 4°C. Counterstains of DAPI or NeuroTrace™ (for lesion experiment tissues) (ThermoFisher Scientific) were performed after primary antibody staining, at a dilution of 1:1000 and 1:100, respectively. For free-floating sections, immunostaining was performed with the same dilution of primary antibodies, with incubations being 1 day or 3 hours, respectively.

### Slide scanning and anatomical reconstructions

Serially mounted coronal sections were imaged with an AZ100 automated slide scanning microscope equipped with a 4x 0.4-NA objective (Nikon). Image processing and analysis were done using BrainJ as previously described^40^. Briefly, sections were registered using 2D rigid-body registration. Ilastik was used to segment cell bodies and neuronal processes of each section. Representative images were imported and labeled to create a training set of cell bodies, neural processes, and background fluorescence for the algorithm across all images, which then created probabilistic assignment of detected features. The resulting probability masks and cell coordinates were aligned to the Allen Brain Reference Atlas (ABA) using Elastix. These aligned cell counts were then used to delineate the forelimb-specific subregions of Pf and VAL in the ABA. GRIN lens placements in imaging animals were manually assessed by matching the ABA, updated with the forelimb-related Pf and VAL, with coronal sections of stained tissue and marking the center lenses (**Figure S3A/B)**. Lesioned areas were manually delineated from NeuroTrace staining on all sections and registered and aligned to the updated ABA. Lesion volumes were visualized using BrainRender^63^. The percentage of ABA areas affected by the lesions were quantified using custom MATLAB code.

### Pre-learning cell ablation and behavioral training

Vglut2-Cre mice (n=18) and WT littermates (n=20) were used for all ablation experiments. Pf or VAL were bilaterally targeted AAV-flex-taCasp3-TEVp (AAV5), which triggers cell death through apoptosis^43^. All mice were injected with the flexed-taCasp3 virus, resulting in expression of caspase3 only in Vglut2-Cre mice. Stereotaxic surgeries with viral injections and headpost implantation took place as described above, with bilateral craniotomies and injections targeting Pf or VAL. Surgeries took place 3 weeks before STT training began. Mice were trained in the STT in block 1 until they reached high performance criteria (three consecutive day average of 65% hit ratio) or for a maximum 35 days, whichever came first. Mice were trained in block 2 until reaching high performance criteria, or for a maximum 19 days.

### Post-learning ablation and behavioral training

Post-learning lesion mice underwent surgeries as described above. Vglut2-Cre mice (n =11) and WT littermates (n=10) were used. The initial surgeries for headbar implantation took place one week before STT training began. All mice were trained in block 1 for 28 days, and had their second surgery of AAV-flex-taCasp3-TEVp (AAV5) injection the following day. Mice were given ad lib food the day before the surgery, and for three days post-surgery. After that, they returned to a food restriction regimen for three weeks, where they maintained 80-85% of their pre-surgery weight, to allow for the expression of the virus. Mice were then trained on the same target as block 1 for 9 consecutive test days.

### Statistical analysis

Statistical tests were performed with GraphPad Prism9.

## Acknowledgements

We would like to thank Luke Hammond and Humberto Avila, and Dr. Darcy Peterka from the Zuckerman Institute’s Cellular Imaging Platform for guidance with analysis of the lesion volumes in Imaris and with BrainJ. We would also like to thank Dr. Darcy Peterka for his support and guidance when collecting 2-photon calcium imaging data. We would like to thank Ramin Khajeh for discussions regarding neural analysis. We would like to thank Gabriela Martins, Mariana Correia, and Drew Baughman for additional support through lab and mouse colony management.

## Funding

National Institutes of Health (NINDS) F31NS111853 (L.J.S.) National Institutes of Health BRAIN Initiative (NINDS) K99NS126307 (A.C.M.)

National Institutes of Health (NINDS) K99NS128250 (V.R.A.) National Institutes of Health (NIMH) F32MH118714 (V.R.A.) National Institutes of Health (NINDS) R00NS114194 (J.M.M.) National Institutes of Health BRAIN Initiative (NINDS) U19NS104649 (R.M.C.)

Swiss National Science Foundation Postdoc fellowship P2EZP3_172128 (A.C.M.)

Swiss National Science Foundation Postdoc fellowship P400PM_183904 (A.C.M.)

Simons-Emory International Consortium on Motor Control (R.M.C., A.C.M.)

## Author contributions

L.J.S. designed the study, analyses, and wrote the manuscript, in collaboration with A.C.M. and R.M.C. L.J.S. and A.C.M. made figures and illustrations. L.J.S. performed surgeries. L.J.S. performed imaging experiments. L.J.S. and T.X.C. performed lesion experiments. A.C.M. and T.X.C. wrote task control code. L.J.S. and T.X.C. processed and analyzed brain tissue. L.J.S. and ACM analyzed mouse behavior data. L.J.S., A.C.M., and V.R.A. analyzed 2-photon calcium imaging data. J.M.M. advised on analysis of 2-photon calcium imaging data. All authors reviewed and edited the manuscript.

## SUPPLEMENTAL FIGURES

**Figure S1.**
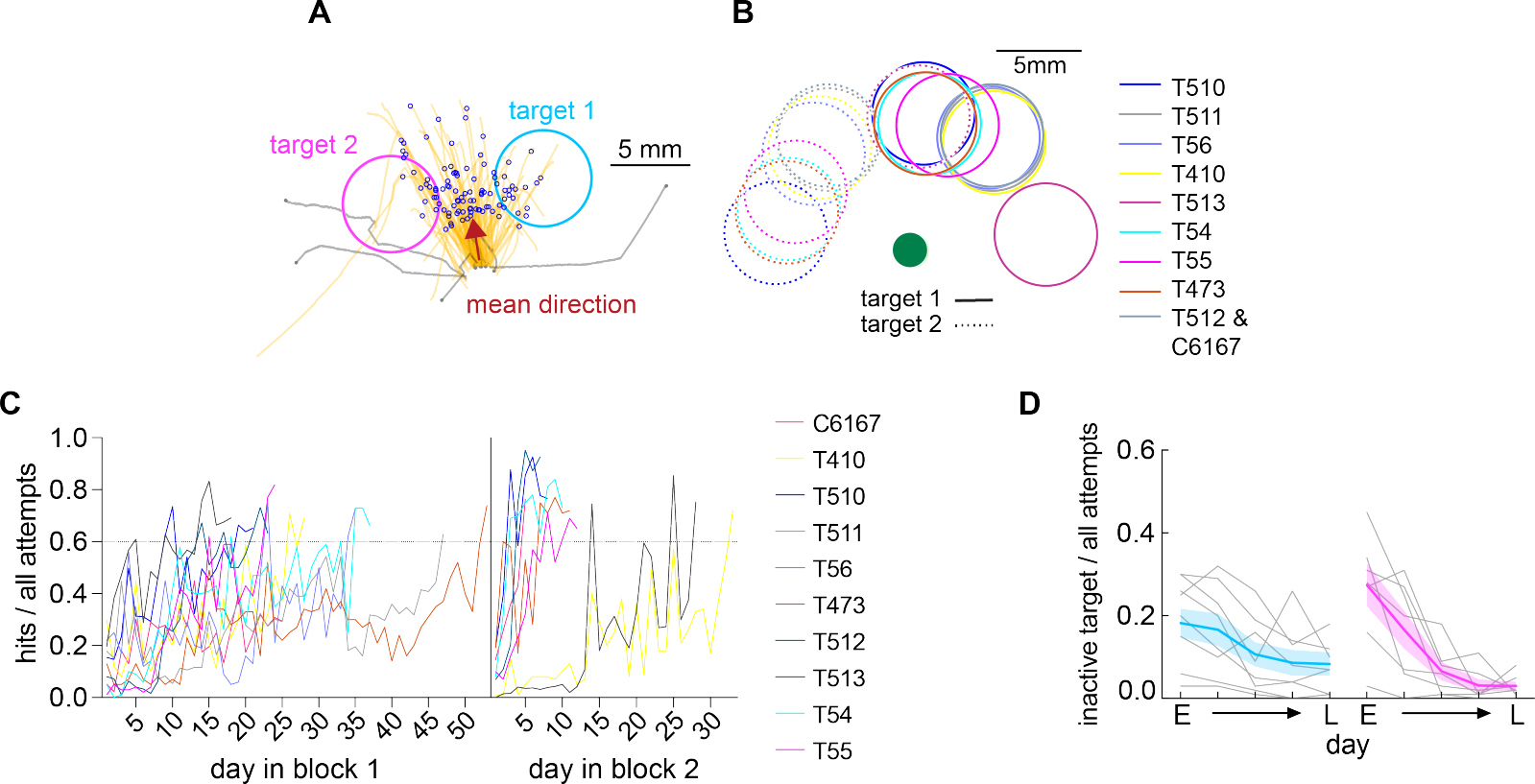
Target selection and behavioral training of spatial target task. (A) Representative behavior of a single animal from last day of pre-training. The red arrow shows the mean hit direction from which targets are defined 40° to the left and right. (B) Individually defined targets for all animals plotted relative to start position (filled green circle). Target 1 for block 1 in solid circle and target 2 for block 2 in dashed circle. (C) Hit ratio (# hits/ all attempts) for all animals across two blocks of training under 2-photon microscope. Individual animals plotted with separate colored lines. (D) Inactive target hit ratio (Mixed-effects model: day in block: *p < 0.001) Mean ± SEM is shown in thick lines with shaded bounds (block 1: blue, block 2: magenta), single animals shown in gray lines (n = 10).

**Figure S2.**
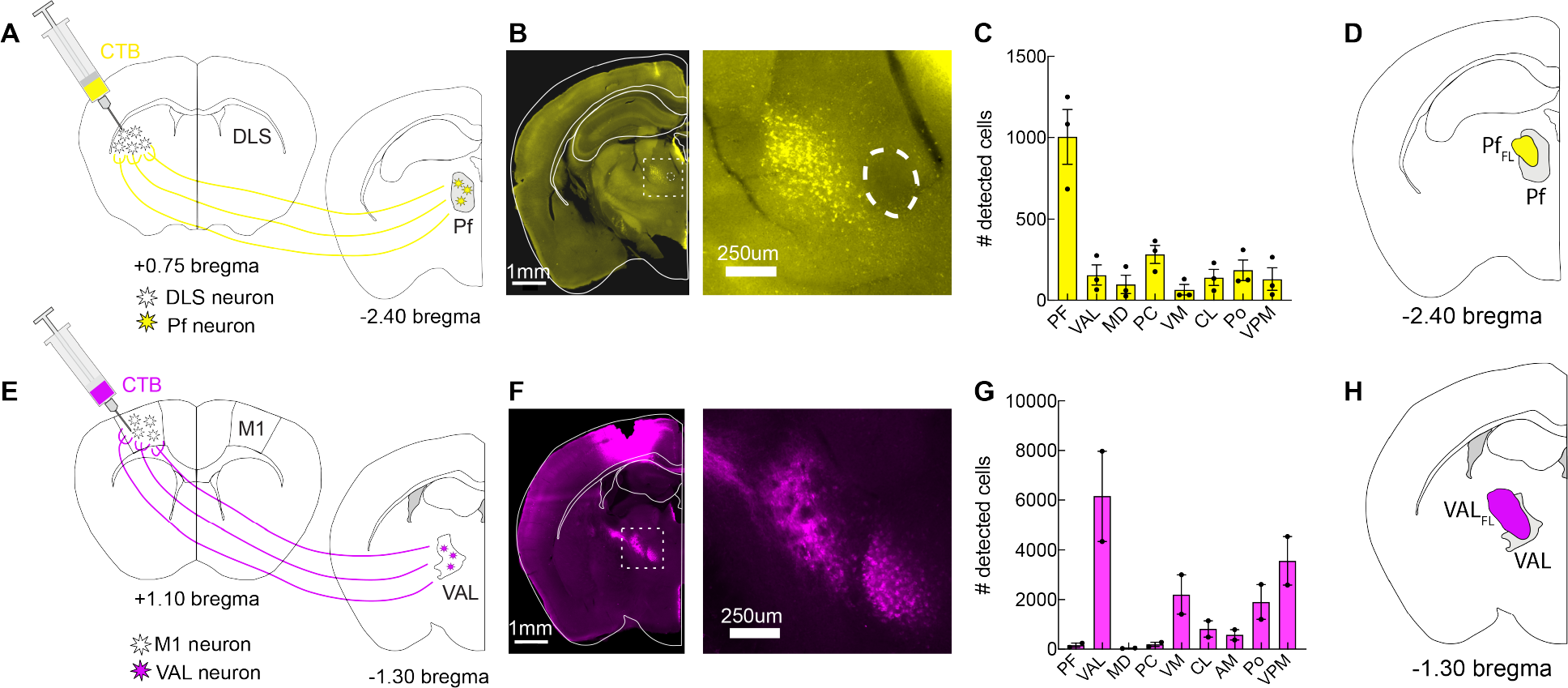
Pf and VAL are the dominant forelimb related thalamic nuclei. (A)Schematic of experimental design showing a coronal ABA section at +0.75 mm bregma and CTB injection (yellow) into forelimb DLS and at -2.40 mm bregma showing the center of Pf. (B)Left: representative section at center of Pf with ABA overlay. Fasciculus retroflexes market with dotted circular outline, and used as an anatomical marker for Pf. Right: magnified image of inset from dashed outline. (C) Number of detected cells in thalamic nuclei after CTB injection into forelimb DLS (n = 3 mice). (D) Coronal ABA section at -2.40 bregma highlighting the Pf (gray) with Pf_FL_ defined from CTB tracing overlayed (yellow) (E)as in (A), but with CTB (magenta) injection in the caudal forelimb area of M1 and ABA sections at the level of M1 and VAL. (F) Left: representative section at center of VAL with ABA overlay. Right: inset as indicated in dashed outline. (G) As in (C), but after CTB injection into CFA of M1 (n = 2 mice). (H) Coronal ABA section at -1.30 bregma highlighting VAL (gray) with VAL_FL_ defined from CTB tracing overlayed (magenta) (E and F) Mean ± SEM shown. Acronyms: PF, parafascicular nucleus; VAL, ventroanterior/ventrolateral nuclei; MD, mediodorsal nucleus; PC, paracentral nucleus; VM, ventromedial nucleus; CL, central lateral nucleus; AV, anteroventral nucleus; AM, anteromedial nucleus; Po, posterior complex of the thalamus; VPM, ventral posteromedial nucleus.

**Figure S3.**
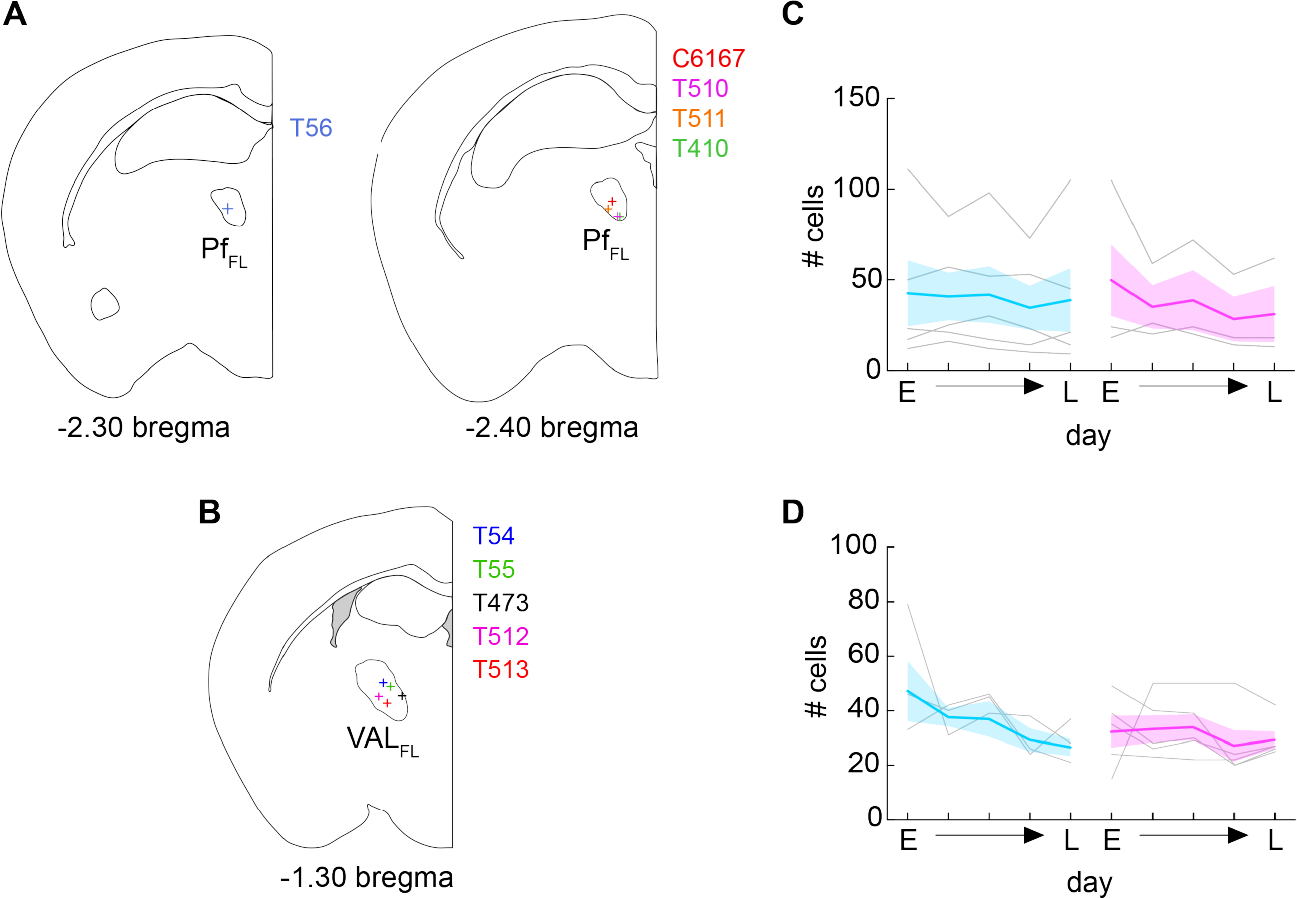
GRIN lens locations and number of extracted cells over training of spatial target task. (A)Center of GRIN lens implants for PF imaging animals (+) (n = 5 mice) in coronal plane. Left: coronal plane at -2.30 mm bregma. Right: coronal plane at -2.40 mm bregma. PF_FL_ indicated in outlined section. (B) Center of GRIN lens implants of VAL imaging animals (+) (n = 5 mice) in coronal plane at -1.30 mm bregma. VAL_FL_ indicated in outlined section. (C) Number of cells on five days of imaging over block 1(blue) and block 2 (magenta) of STT training. Early (E) day, late (L) day, and 3 equally spaced days over each block. (D) As in (C), but for VAL animals. (C-D) Mean ± SEM depicted with thick colored lines and shaded bounds. Individual mice in gray lines.

**Figure S4.**
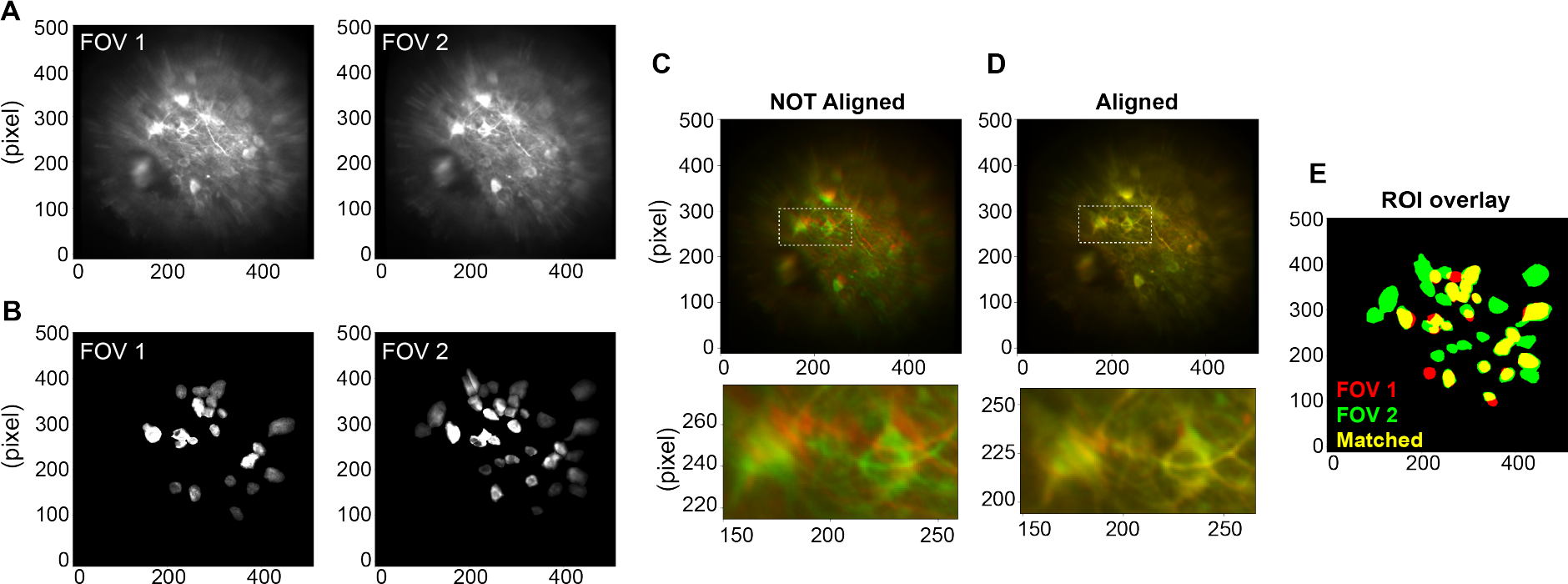
Tracking cells over multiple days of 2-photon imaging. (A)Average projections of example imaging FOV over two days. FOV1 from the late day of block 1 and FOV2 from the early day of block 2. (B) Max projections of detected ROIs from the imaging FOV1 and FOV2 as seen in (A). (C) Top: Representative overlay of FOV1 and FOV2 before non-rigid alignment; late day of block 1 (red) and early day of block 2 (green). White dotted line indicating inset. Bottom: Inset area before alignment (D) Top: Representative alignment of FOV1 and FOV2 after non-rigid alignment; late day of block 1 (red) and early day of block 2 (green). White dotted line indicating inset. Bottom: Inset area after alignment. (E) Overlay of detected ROIs from FOV 1 (red), FOV 2 (green) and matched overlapping areas (yellow) from both sessions.

**Figure S5.**
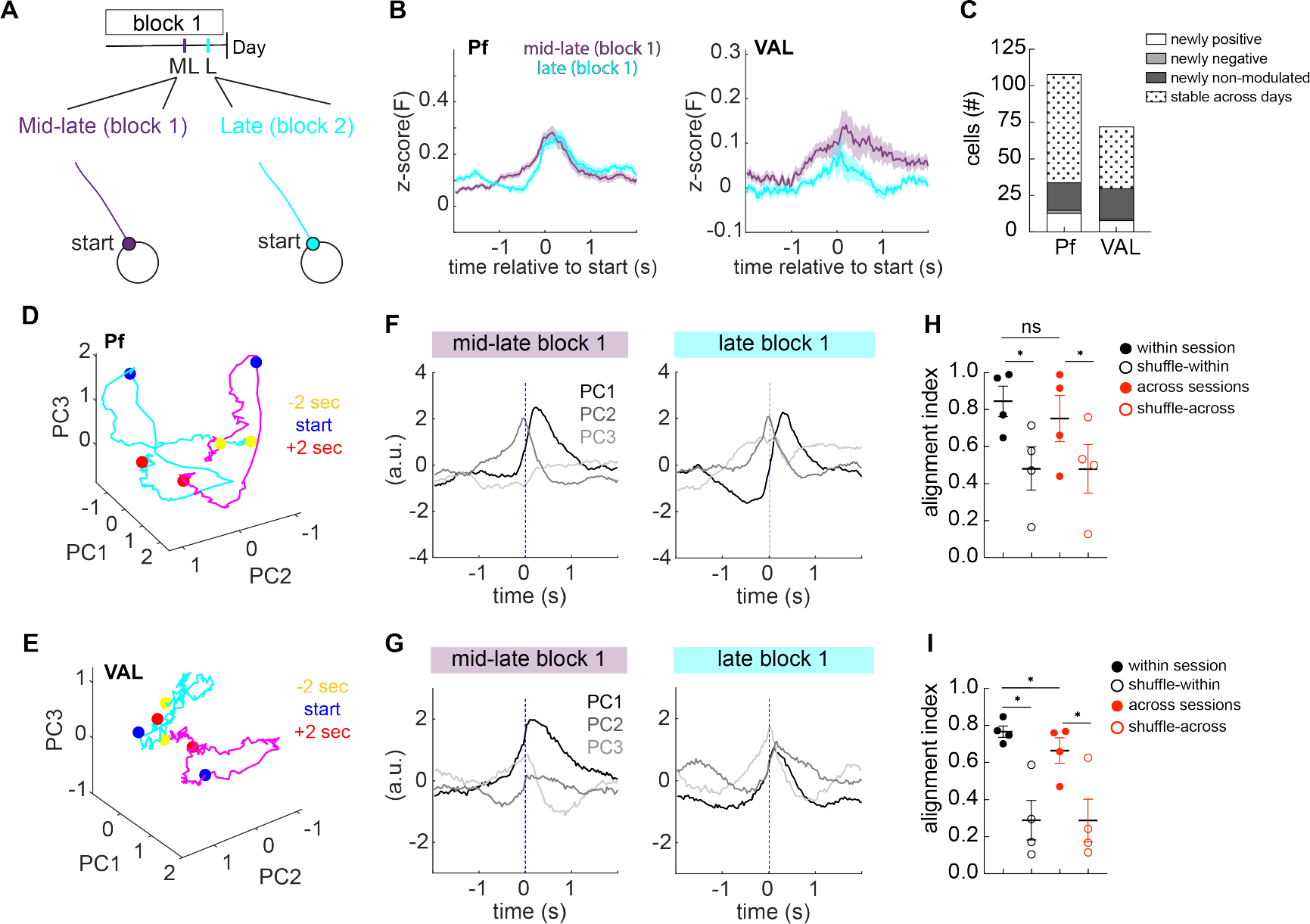
Low dimensional neural activity alignment unchanged between two days with similar behavior. (A) Schematic showing the selected days for cell matching; mid-late day of block 1 (ML) and late day of block 1 (L) and depicting the relative behavioral window around movement start. (B) Left: trial-averaged fluorescence aligned to movement start for matched cells on mid-late block 1 (purple) and late block 1 (cyan) for Pf_FL_ (n =108 cells, 4 mice; Wilcoxon matched-pairs signed rank test: p = 0.722). Right: same for VAL_FL_ (n = 72 cells, 4 mice; Wilcoxon matched-pairs signed rank test: p = 0.052) Mean ± SEM (thick colored line with shaded bounds). (C) For Pf_FL_ and VAL_FL_ matched cells, cellular responses on the late day of block 1, tracking each cell’s response relative to the response it had on the mid-late day of block 1. Cells either became positively modulated (white), negatively modulated (light gray), non-modulated (dark gray), or had a stable modulation on both days (black dots). (D) Neural activity of Pf_FL_ around movement start projected into top 3 PC space for mid-late day of block 1 (magenta trace) and late day for block 1 (cyan trace). -2 seconds before movement onset (green circle), movement start (blue circle), and +2 seconds after movement start (red circle). (E) Same as (D), for VAL_FL_ matched neuronal populations. (F) Same neural activity as in (D) projected onto the top 3 PCs and plotted against time and aligned to movement start (dashed blue line). (G) Same as (F), for VAL_FL_ matched neuronal populations. (H) For Pf_FL_, average alignment of subspaces occupied by top 3 PCs within (filled red) and across (filled black) sessions with matched cells (same sessions D-E). Alignment within sessions is not significantly different than across, but is higher than within-shuffle (open black) (by 2-way repeated measures ANOVA: Day of alignment: p = 0.217; shuffle: *p = 0.008; Day of alignment x shuffle: p = 0.148; Šídák’s multiple comparison: within vs across, p = 0.131 ; within vs within-shuffle, *p = 0.003). Alignment between across and across-shuffle (open red) are also significantly different (*p = 0.008 Šídák’s multiple comparison, across vs shuffle-across). (I) Same as (H), for VAL_FL_ matched neuronal populations. Alignment within sessions is significantly different than across in post-hoc analysis, and is higher than within-shuffle (by 2-way repeated measures ANOVA: Day of alignment: p = 0.232; shuffle: *p = 0.035; Day of alignment x shuffle: p = 0.058; Šídák’s multiple comparison: within vs across, *p = 0.045; within vs within-shuffle, *p<0.001). Alignment between across and across-shuffle (open red) are also significantly different (*p = 0.001 Šídák’s multiple comparison, across vs shuffle-across).

**Figure S6.**
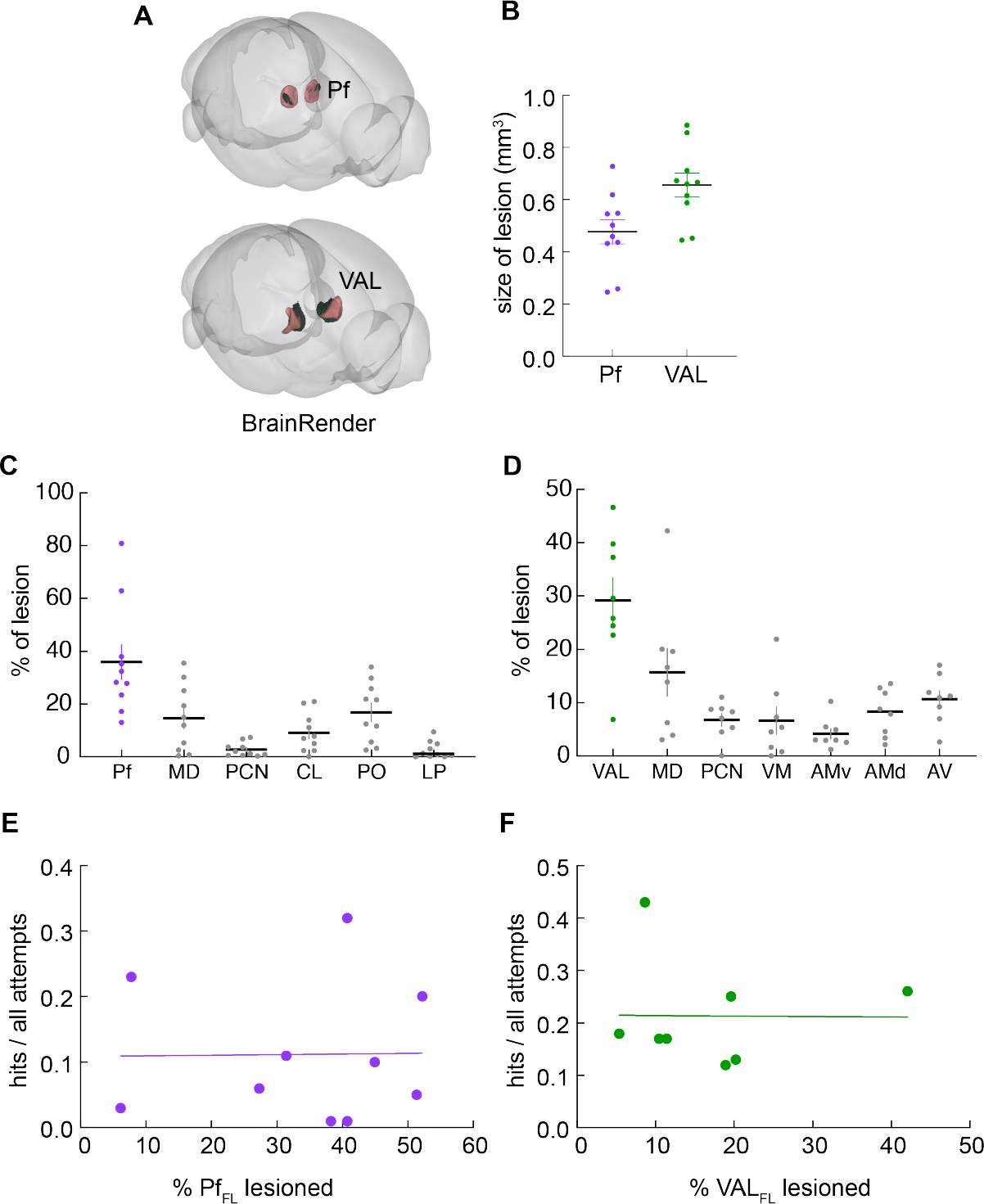
Size of pre-learning lesion size does not correlate with performance. (A) Representative 3D reconstruction of lesions areas (green-volume) of bilateral Pf (top – red volume) and VAL (bottom – red volume) lesion. Made using BrainJ outputs and BrainRender for visualization. (B) Total lesion volume (mm^3^) for Pf_pre-lesion_ (purple) and VAL_pre-lesion_ (green) mice. (C) Relative percent of total lesion in thalamic nuclei for Pf_pre-lesion_ mice. Relative percent of lesion in Pf (purple), other thalamic areas (gray). (D) Same as in (D) but for VAL_pre-lesion_ mice. Relative percent of lesion in VAL (green), other thalamic areas (gray) (E) Percent of Pf lesioned in Pf_pre-lesion_ miced versus hit ratio on the last day of block 1 (hits/all attempts). n=10 mice. Fitted regression line(purple line) shows there is no significant correlation between percent of Pf_FL_ lesioned and hit ratio. (Simple linear regression, slope different from zero: p=0.968). (F) Same as (E), but for VAL_pre-lesion_ lesion cohort, n=8. Fitted regression line (green) shows there is no significant correlation between percent VAL_FL_ lesioned and hit ratio. (Simple linear regression, slope different from zero: p=0.982). (B-D) Mean ± SEM shown. Pf_pre-lesion_ n=10 mice, VAL_pre-lesion_ n=8 mice. (C and D) Acronyms: MD: mediodorsal nucleus; PCN: paracentral nucleus; CL: central lateral; PO: posterior complex of the thalamus; LP: lateral posterior nucleus; VM: ventral medial nucleus; AV: Anteroventral nucleus; AMv: Anteromedial nucleus, ventral part; AMd: Anteromedial nucleus, dorsal part.

**Figure S7.**
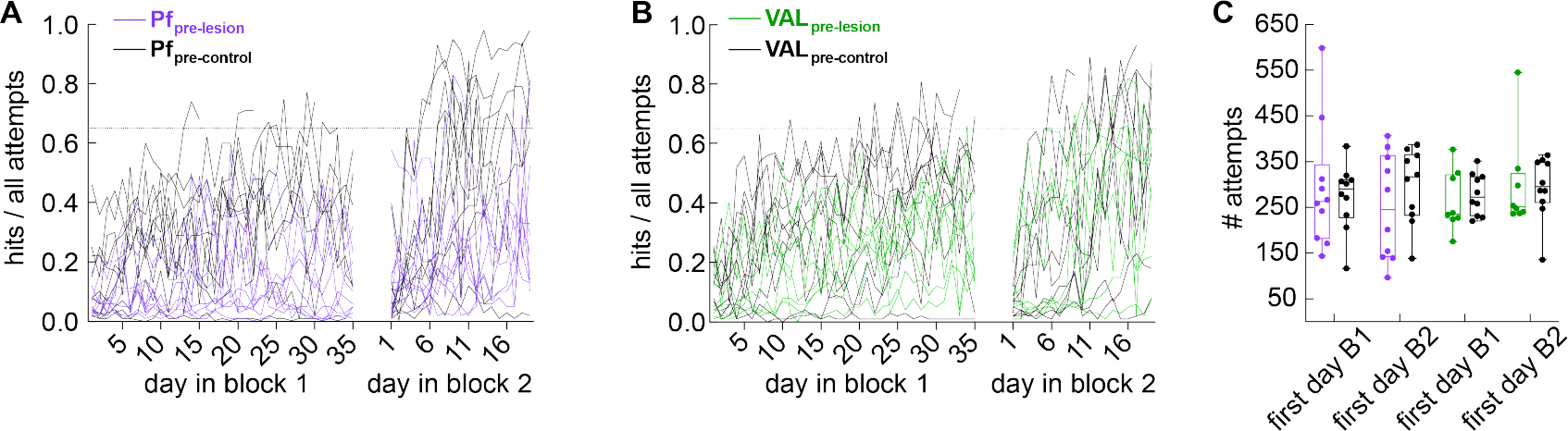
Behavioral training of pre-learning Pf and VAL lesioned mice in spatial target task. (A) Hit ratio (hits / all attempts) of each animal in pre-learning lesion experiment. High performance criterion (65% hit ratio) indicated in dotted black line. Pf_pre-lesion_ (purple), n = 10; Pf_pre-control_ (black lines), n = 10. (B) Same as (A) but for VAL pre-learning lesion experiment. VAL_pre-lesion_ (green lines), n = 8; Pf_pre-control_(black lines), n = 10. (C) Engagement (# of attempts) on the first day of block 1 and block 2. There is no difference in level of engagement for all control and lesion groups. (unpaired t-test: block 1 Pf: p = 0.713; block 2 Pf: p = 0.318; block 1 VAL: p = 0.606; block 2 VAL: p = 0.895). Data shown as box and whisker plot, whiskers to the min and max values.

**Figure S8.**
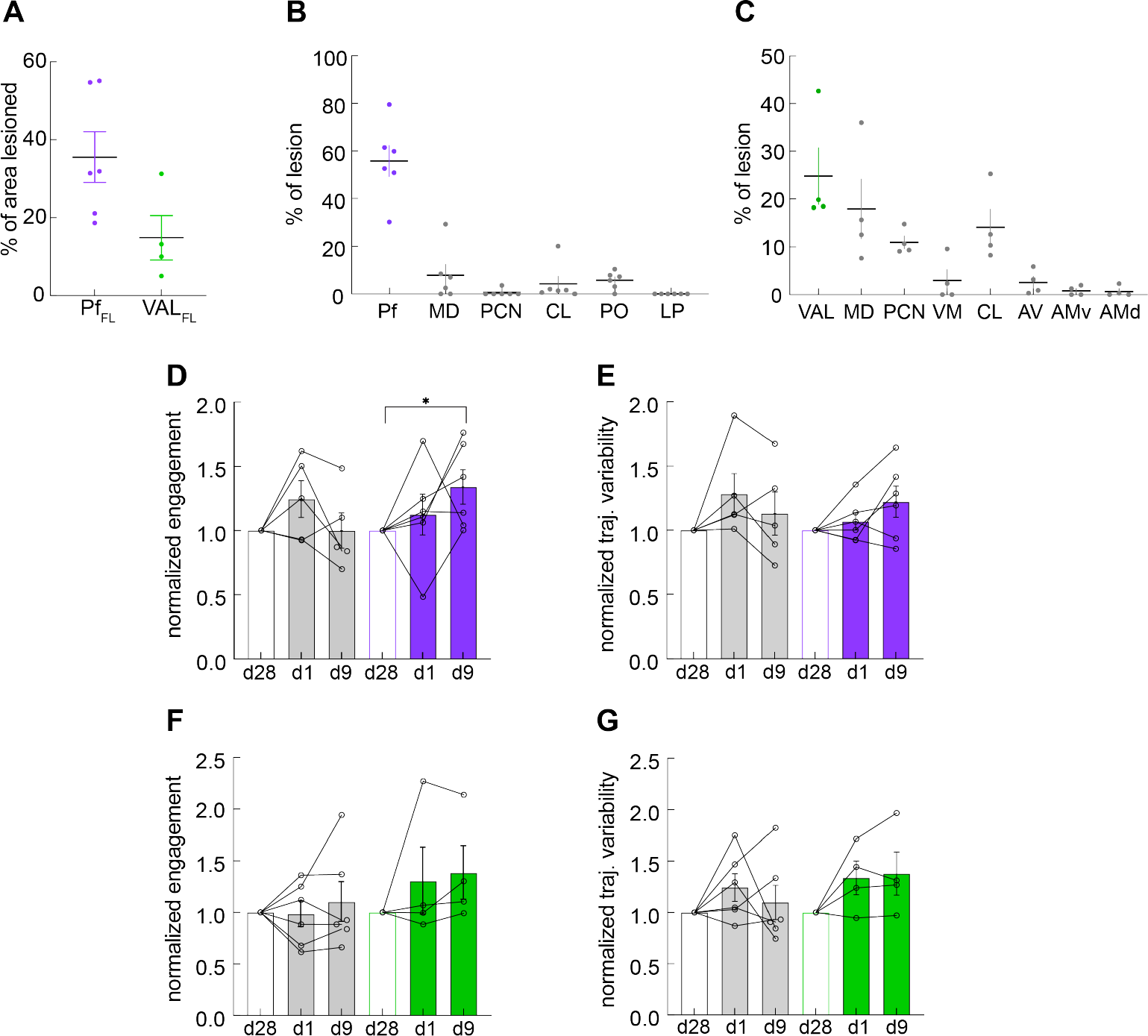
Behavioral training of post-learning Pf and VAL lesioned mice in spatial target task. (A)Percent of Pf_FL_ (purple) and VAL_FL_ (green) in post-learning lesion groups. (B) Relative percent of lesion in Pf (purple) and neighboring thalamic nuclei (gray). (C) Relative percent of lesion in VAL (green) and neighboring thalamic nuclei (gray) (D) normalized engagement (# of attempts) of Pf groups on d1 and d9 of post-lesion test. Pf_post-lesion_ group increases number of attempts over 9 post-lesion test days. (Wilcoxon test for d9 vs d28: Pf_post-lesion_, *p=0.031; Pf_post-control_, p=0.813) (E) normalized variability along average trajectory (std in mm) for Pf_post-lesion_ (purple) and Pf_post-control_ (gray). (Wilcoxon test for d9 vs d28: Pf_post-lesion_, p=0.124; Pf_post-lesion_, p>0.999) (F) Same as (D), for VAL_post-lesion_ (green) and VAL_post-control_ (gray). (Wilcoxon test for d9 vs d28: VAL_post-lesion_, p=0.250; VAL_post-control_, p>0.999) (G) Same as (E), for VAL groups. There is no effect for VAL_post-lesion_ group. (Wilcoxon test for d9 vs d28: VAL_post-lesion_, p=0.250) (D-G) performance metrics on the first (d1) and last (d9) day of post-lesion test. Data normalized to the values on the last day of training before lesion (d28, open white bars). Filled colored bars denote post-lesion days for Pf_post-lesion_ (purple), VAL_post-lesion_ (green), and control groups (gray). Individual animal data plotted with open circles. (A-G) Mean ± SEM shown.1: group: p = 0.216. Block 2: group: p = 0.090). (N) Same as in (M), for VAL lesion groups. VAL_pre-lesion_ mice overshoot the target more than VAL_pre-control_ animals in block 1 (2-way repeated measures ANOVA: Block 1: group: *p = 0.019. Block 2: group: p = 0.293). Šídák’s multiple comparison test VAL_pre-lesion_ vs VAL_pre-control_ late block 1: *p = 0.023.

